# Matrix mechanotransduction mediated by thrombospondin-1/integrin/YAP signaling pathway in the remodeling of blood vessels

**DOI:** 10.1101/814533

**Authors:** Yoshito Yamashiro, Bui Quoc Thang, Karina Ramirez, Seung Jae Shin, Tomohiro Kohata, Shigeaki Ohata, Tram Anh Vu Nguyen, Sumio Ohtsuki, Kazuaki Nagayama, Hiromi Yanagisawa

## Abstract

**Rationale:** The extracellular matrix (ECM) initiates mechanical cues and transduces intracellular signaling through matrix-cell interactions. In the blood vessels, additional mechanical cues derived from the pulsatile blood flow and pressure play a pivotal role in homeostasis and disease development. Currently, the nature of the cues from ECM and how they coordinate with a mechanical microenvironment in large blood vessels to maintain the integrity of the vessel wall are not fully understood.

**Objective:** The aim of this study was to elucidate the crucial mediator(s) and molecular signaling pathway(s) involved in matrix mechanotransduction during remodeling of the vessel wall.

**Methods and Results:** We performed secretome analysis using rat vascular smooth muscle cells (SMCs) under cyclic stretch and examined matrix-cell interactions and cell behavior. We found that the matricellular protein thrombospondin-1 (Thbs1) was secreted upon cyclic stretch and bound to integrin αvβ1, thereby recruiting vinculin and establishing focal adhesions. RNA-sequence (RNA-seq) analysis revealed that deletion of Thbs1 in vitro markedly affected the target gene expression of Yes-associated protein (YAP). Consistently, we found that Thbs1 promotes nuclear shuttling of YAP in response to cyclic stretch, which depends on the small GTPase Rap2 and Hippo pathway, and is not influenced by alteration of actin fibers. Deletion of *Thbs1* in mice inhibited Thbs1/integrin/YAP signaling, leading to maladaptive remodeling of the aorta in response to pressure overload by transverse aortic constriction (TAC), whereas it suppressed neointima formation upon carotid artery ligation, exerting context-dependent effects on the vessel wall.

**Conclusions:** Thbs1 serves as a mechanical stress-triggered extracellular mediator of mechanotransduction that acts via integrin αvβ1 to establish focal adhesions and promotes nuclear shuttling of YAP. We thus propose a novel mechanism of matrix mechanotransduction centered on Thbs1, connecting mechanical stimuli to YAP signaling during vascular remodeling in vivo.

**Subject codes:**

- Vascular Disease
- Genetically Altered and Transgenic Models
- Vascular Biology
- Cell Signaling/Signal Transduction

## Introduction

The extracellular matrix (ECM) is fundamental to cellular and tissue structural integrity and provides mechanical cues to initiate diverse biological functions^1^. The quality and quantity of ECM determine tissue stiffness and control gene expression, cell fate, and cell cycle progression in various cell types^2, 3^. ECM-cell interactions are mediated by focal adhesions (FAs), the main hub for mechanotransduction, connecting ECM, integrins and cytoskeleton^1, 4^. In the blood vessels, an additional layer of mechanical cues derived from the pulsatile blood flow and pressure play a pivotal role in homeostasis and disease development. However, how cells sense and integrate different mechanical cues to maintain the blood vessel wall is largely unknown.

Hippo effector Yes-associated protein (YAP) and transcriptional coactivator with PDZ-binding motif (TAZ) serve as an on-off mechanosensing switch for ECM stiffness^5^. YAP activity is regulated by a canonical Hippo pathway; MST1/2 (mammalian sterile 20-like 1/2 kinases) and their downstream kinases LATS1/2 (large tumor suppressor 1/2), through which YAP is phosphorylated (inactivate) and retained in cytoplasm by binding to 14-3-3 protein^6^. YAP activity is also regulated by Wnt/β-Catenin and G-protein-coupled receptor (GPCR) signaling pathways^7, 8^. However, regulation of YAP activity is highly context-dependent and extracellular regulators are not fully understood^6, 9^.

Thrombospondin-1 (Thbs1) is a homotrimeric glycoprotein with a complex multidomain structure capable of interacting with a variety of receptors such as integrins, cluster of differentiation (CD) 36 and CD47^10^. Thbs1 is highly expressed during development and reactivated in response to injury^11^, exerting domain-specific and cell type-specific effects on cellular functions. We have recently shown that Thbs1 is induced by mechanical stretch and is a driver for thoracic aortic aneurysm in mice^12, 13^.

In the current study, we identified Thbs1 as an extracellular regulator of YAP, which is induced by mechanical stress and acts through FAs independent of actin remodeling. Mechanistically, Thbs1 binds integrin αvβ1 and strengthen FAs, aiding in translocation of YAP to the nucleus. Deletion of Thbs1 in vivo exhibited altered vascular remodeling in response to pressure overload and flow cessation. Taken together, Thbs1 mediates dynamic interactions between mechanical stress and the YAP-mediated transcriptional cascade in the blood vessel wall.

## Methods

Detailed descriptions are provided in the Online Data Supplement.

### Cell culture and reagents

Rat vascular SMCs (Lonza, R-ASM-580) from the aorta of 150-200 grams adult male Sprague-Dawley rats, were grown in DMEM with 20% (v/v) fetal bovine serum and 1 x Antibiotic-Antimycotic (Thermo Fisher Scientific). HUVECs were grown in HuMedia-MvG (KURABO) supplemented with 5% (v/v) fetal bovine serum and growth factors. The cell lines were used as passage 7 to 12 and tested for mycoplasma contamination using Mycoplasma detection set (TaKaRa, #6601) following the manufacture’s protocol (Online Fig. XIX). Brefeldin A (B5936) was purchased from Sigma-Aldrich. PF-00562271, FAK inhibitor (S2672) was purchased from Selleck. Cytochalasin D (037-17561) and Latrunculin A (125-04363) were purchased from Wako. Jasplakinolide (AG-CN2-0037-C050) was purchased from AdipoGen Life Sciences. Recombinant human thrombospondin-1 was purchased from R&D Systems (3074-TH-050).

### In vitro mechanical stretch

Cyclic stretch was performed using a uniaxial cell stretch system (Central Workshop Tsukuba University) following 24 h of serum starvation as described previously^13^. Rat vascular SMCs or HUVECs were plated (3 × 10^4^ cells) on silicon elastomer bottomed culture plates (SC4Ha, Menicon Life Science) coated with cell attachment factor containing gelatin (S006100, Thermo Fisher Scientific) and subjected to cyclic stretch with a frequency of 1.0 Hz (60 cycles/min) and 20% strain for the indicated time.

### Mice

C57BL/6J (wild-type) mice were purchased from Charles River, Japan and *Thbs1* null mice were purchased from Jackson Laboratory (B6.129S2-*Thbs1*^*tm1Hyn*^/J) and crossed with C57BL/6 mice for line maintenance. Male mice were used for the experiments and mice were kept on a 12 h/12 h light/dark cycle under specific pathogen free condition and all animal protocols were approved by the Institutional Animal Experiment Committee of the University of Tsukuba.

### Statistical analysis

All experiments are presented as means ± standard error of the mean (SEM). Statistical analysis was performed using Prism 7 (Graph Pad). *P* < 0.05 denotes statistical significance.

### Data availability

The acquisition of online accession numbers for the secretome data is jPOST ID: JPST000600/PXD013915 (http://jpostdb.org) and for the RNA-seq data is GSE131750 (GEO accession). All data supporting the findings of this study are available from the corresponding authors on reasonable request.

## Results

### Cyclic stretch induced secretion of ECM and cell adhesion molecules involved in blood vessel development

Vascular ECM is synthesized by SMCs from mid-gestation to the early postnatal period and determines the material and mechanical properties of the blood vessels^14^. Owing to the long half-life of ECM, blood vessels are generally thought to stand life-long mechanical stress without regeneration, particularly in large arteries. However, it is not completely known whether local micro-remodeling system exists for the maintenance of the vessel wall. To search for a potential remodeling factor(s), we first performed cyclic stretch experiments (1 Hz; 20% strain) using rat vascular SMCs for 20 hours (h) with or without Brefeldin A (BFA), a protein transporter inhibitor. Conditioned media (CM) were analyzed using quantitative mass spectrometry and 87 secreted proteins with more than two-fold differences in response to cyclic stretch were identified (Fig. 1A-B, Online Figs. I and II). Gene ontology (GO) enrichment analysis using GO Consortium (http://www.geneontology.org) revealed that proteins involved in ECM organization and structural constituents were highly enriched (Fig. 1C). Ingenuity Pathway Analysis (IPA) was conducted for biological interaction network among proteins in ECM-cell adhesions and blood vessel development (Fig. 1D) and Thbs1 showed multiple interactions in both categories. We confirmed the increased expression and secretion of Thbs1 after cyclic stretch (Fig. 1E), suggesting that Thbs1 may play an important role in response to dynamic vessel microenvironments.

**Figure 1.**
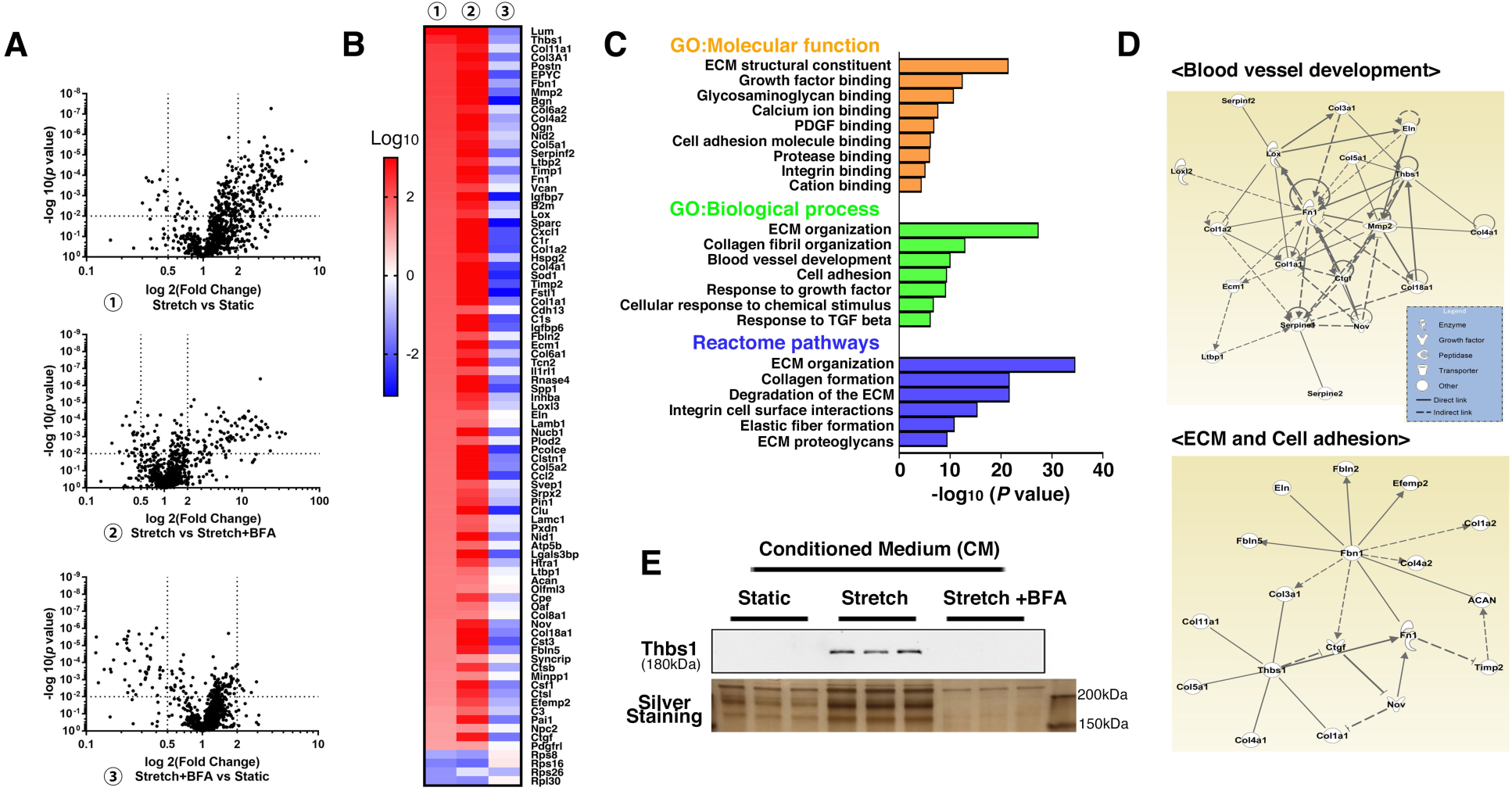
Comprehensive secretome analysis of smooth muscle cells under cyclic stretch. **A**, Volcano plots of protein expressions between the two conditions (1: Stretch vs Static, 2: Stretch vs Stretch+BFA, 3: Stretch+BFA vs Static). Brefeldin A (BFA, 1µM) was used to inhibit secretion. The p-values in the unpaired Student’s t test were calculated and plotted against the fold-change for all identified proteins. Each dot represents mean values (n=3). **B**, Heat map of secreted proteins induced by cyclic stretch. A list of all the proteins is provided in Online Fig. II. **C**, Functional enrichment analysis of 87 proteins, the negative log_10_ of the *P* value. Top enriched GO terms associated with molecular function (orange), biological process (green), and reactome pathway analysis (blue) are shown. **D**, Biological network using Ingenuity Pathway Analysis (IPA). Diagram shows the direct (solid lines) and indirect (dashed lines) interactions among proteins reported in blood vessel development (left) and ECM and cell adhesion (right). **E**, Western blot for Thbs1 from conditioned medium (CM) of rat vascular SMCs in static, stretch (1 Hz, 20 % strain, 20h) with or without BFA (n=3). Entire image of silver staining is provided in Online Fig. I-A.

### Secreted thrombospondin-1 binds to integrin αvβ1 under cyclic stretch

In response to cyclic stretch, SMCs showed upregulation of Early growth response 1 (Egr1) transcription factor, phosphorylated (p) focal adhesion kinase (FAK), Extracellular signal-regulated kinase (ERK), stress-response mitogen-activated protein kinase (MAP) p38 as previously reported (Online Fig. III-A)^13, 15, 16^. Cyclic stretch altered subcellular localization of Thbs1 from peri-nuclear region to the tip of cell on the long axis (Fig. 2A). We examined Thbs1 localization and re-orientation of SMCs as a function of time (Online Figs. III-B, C). SMCs were randomly orientated after 30 min of cyclic stretch but most cells were aligned in a perpendicular position to the stretch direction at 3 h and completely re-orientated by 6 h. Thbs1 localization clearly changed from 3 h after cyclic stretch, which coincided with the re-orientation of actin fibers and elongation of cells (Online Fig. III-D). Since FAs recruit actin stress fibers^17^, we co-stained SMCs with Thbs1 and p-Paxillin, a FA molecule, under cyclic stretch. Thbs1 co-localized with p-Paxillin independent of cell density after cyclic stretch (Figs. 2B, C). Since Thbs1 has well-characterized cell surface receptors, including integrins β1, β3, αv, and CD47, and CD36 (Fig. 2D^10^), we examined which of these receptors bound to Thbs1 in response to cyclic stretch. Immunostaining showed that Thbs1 co-localized with integrins αv and β1, but not with integrin β3, CD47 or CD36 (Fig. 2E-F). Immunoprecipitation of cell lysate obtained from static or cyclic stretch condition with anti-Thbs1 antibody followed by Western blotting confirmed a direct binding between Thbs1 and integrins αv and β1 in the cyclic stretch condition (Fig. 2G). Consistently, Proximity Ligation Assay (PLA) showed that integrins αv and β1 interacted with Thbs1 only in the cyclic stretch condition (Fig. 2H). These data demonstrated that Thbs1 was secreted in response to cyclic stretch and bound to integrin αvβ1.

**Figure 2.**
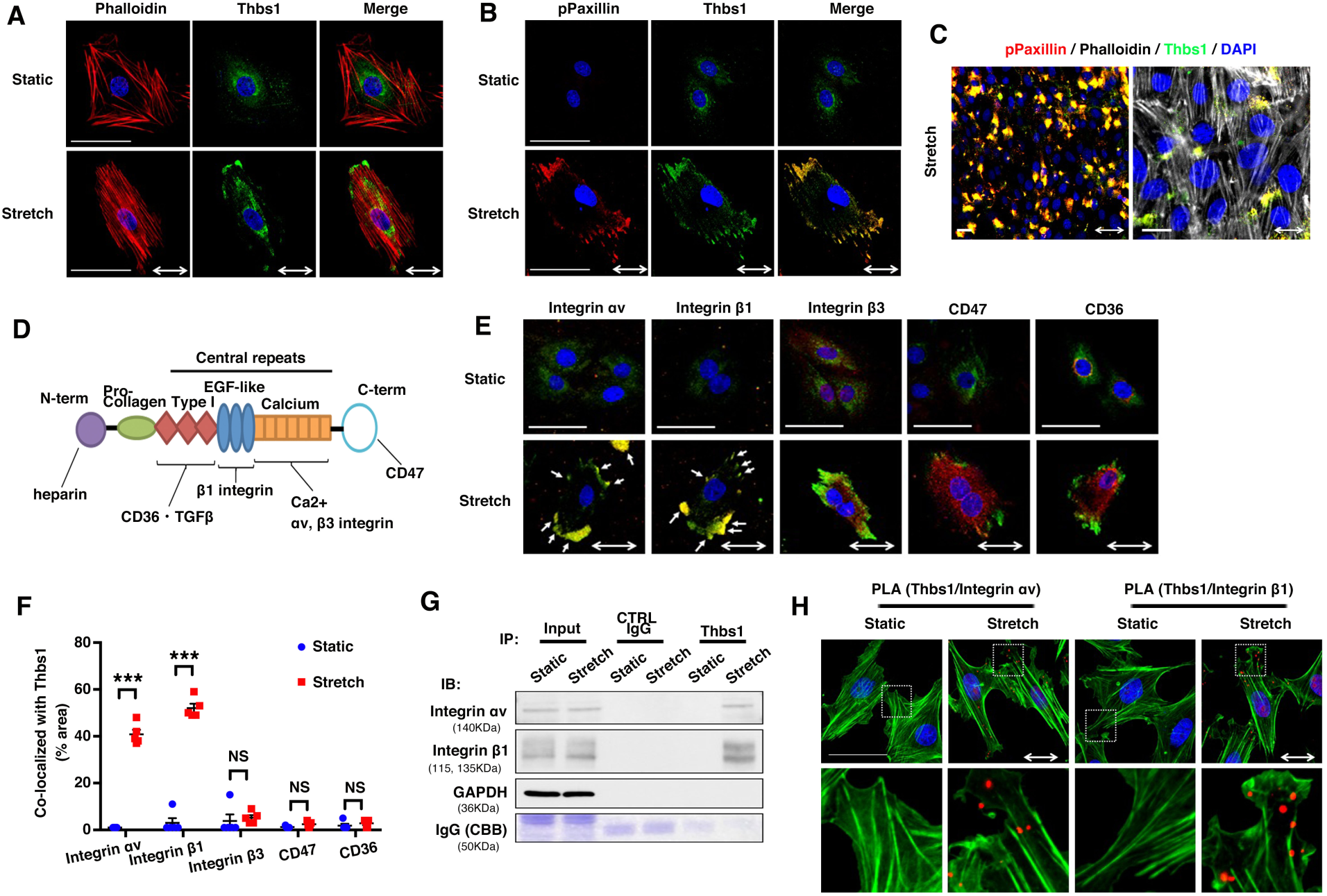
Thbs1 localizes at focal adhesions (FAs) and binds integrin αvβ1 under cyclic stretch. **A**, Cyclic stretch alters localization of Thbs1 (n=5). Representative immunostaining with Phalloidin (red), Thbs1 (green), and DAPI (blue). **B**, Representative image of Thbs1 co-localizes with phosphorylated (p) Paxillin under cyclic stretch (n=5). Immunostained with pPaxillin (red), Thbs1 (green), and DAPI (blue). **C**, Thbs1 localizes at FAs in confluent conditions (n=3). Immunostained with pPaxillin (red), Thbs1 (green), Phalloidin (grey) and DAPI (blue). **D**, Cartoon shows various binding domains of Thbs1 and its receptors. **E**, Representative immunostaining of Thbs1 co-localizes with Integrin αv, and β1 (white arrows), but not with integrin β3, CD47, or CD36 under cyclic stretch (n=5). Immunostained for indicated antibodies (red), Thbs1 (green), and DAPI (blue). **F**, Quantification of co-localization of Thbs1 with molecules shown in (E) using Imaris colocalization software. Bars are mean ± SEM. \*\*\**P* < 0.001, two-way ANOVA. **G**, Immunoprecipitation with anti-Thbs1 or control IgG followed by Western blotting for Integrin αv and β1 (n=2). CBB shows heavy chain of IgG in IP-lysates. **H**, Proximity Ligation Assay (PLA) shows the clusters of Thbs1/Integrin αv or Thbs1/Integrin β1(red dots). The bottom panels show high-magnified images of the white-dashed box in upper panels. DAPI (blue) and Phalloidin (green). In all experiments, rat vascular SMCs were subjected to cyclic stretch (1.0 Hz, 20 % strain for 20 h (A-C, E) or 8 h (G and H). 80-100 cells were evaluated in each immunostaining in A-C, E and H. Two-way arrows indicate stretch directions. Bars are 50 µm.

### Thrombospondin-1 regulates maturation of FA-actin complex and controls cell stiffness

We next examined the effect of deletion of *Thbs1* on cyclic stretch-induced formation of FA and re-orientation of SMCs. *Thbs1* knockout rat vascular SMCs (*Thbs1KO*) were generated using CRISPR-Cas9 genome editing (Online Fig. IV). We evaluated possible off-target effects and confirmed that the expression of *Cpreb3* or *Tet3* was not altered in *Thbs1KO* cells (Online Fig. V-A, B). No obvious phenotypic changes were observed in *Thbs1KO* cells (Online Fig. V-C, D). *Thbs1KO* cells were subjected to cyclic stretch (1 Hz; 20% strain) for 8 h and were compared with control (CTRL) cells with or without 1 µM of BFA. CTRL cells responded to cyclic stretch and re-oriented perpendicular to the direction of stretch, whereas BFA-treated CTRL cells and *Thbs1KO* cells neither elongated nor aligned to the perpendicular position (Figs. 3A-C). Stretch-induced phosphorylation of Paxillin and upregulation of integrin αv and β1 were not observed in *Thbs1KO* cells (Fig. 3D).

**Figure 3.**
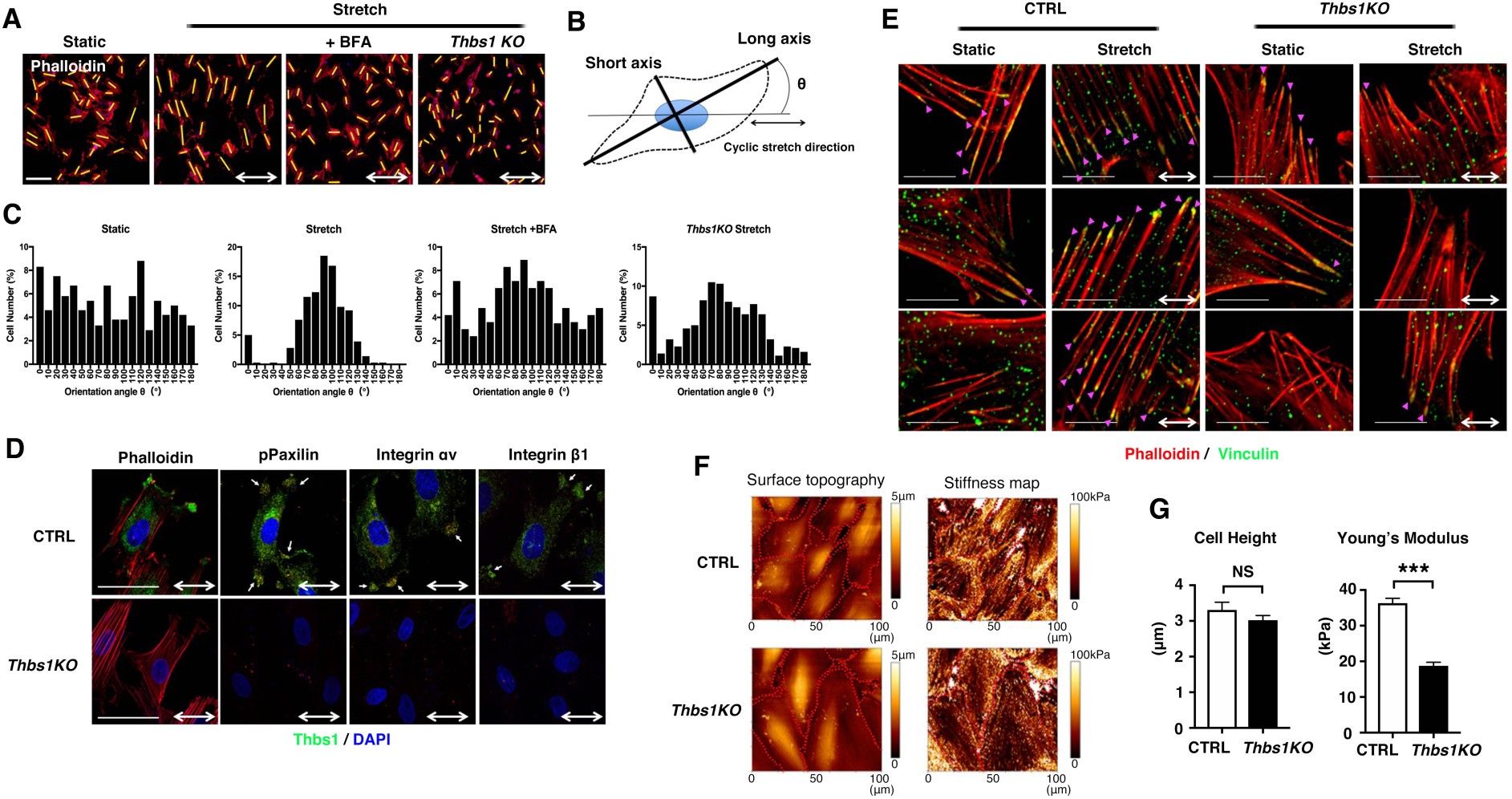
Deletion of Thbs1 affects maturation of focal adhesion and cell stiffness. **A**, Wild-type cells with or without 1 µM of BFA or *Thbs1KO* cells were subjected to cyclic stretch (1.0 Hz, 20% strain, 8h). Two-way arrows indicate the stretch direction. Bars are 50 µm. Phalloidin (red). **B-C**, The orientation of each cell was analyzed by measuring the orientation angle (θ) of the long axis (yellow bars in A) of the ellipse relative to the stretch axis (n=3). **D**, *Thbs1KO* cells show reduced FA formation. Immunostaining with pPaxillin (red), Integrin αv (red), integrin β1 (red), Thbs1 (green), and DAPI (blue) (n=3). Phalloidin (red). White arrows indicate co-localization with Thbs1. Two-way arrows indicate stretch direction. Bars are 50 µm. **E**, Immunostaining of CTRL or *Thbs1KO* cells with vinculin (green) and Phalloidin (red). Rat vascular SMCs were subjected to cyclic stretch (1.0Hz, 20% strain for 8h). Two-way arrows indicate stretch direction. Bars are 25 µm. Arrow heads (purple) show vinculin deposition onto the tip of actin fibers. **F**, Young’s modulus of actin fibers in CTRL (n=129) or *Thbs1KO* (n=86) cells measured using atomic force microscopy. Representative topographic images and stiffness maps are shown. **G**, Cell height and Young’s modulus are shown. Bars are means ± SEM. \*\*\**P* < 0.001, unpaired *t*-test.

Mature FAs are localized at the termini of actin stress fibers and form FA-actin complex^18, 19^. FA-actin complex is stabilized by vinculin deposition and integrin-talin binding, which facilitates PI3K-mediated phosphatidylinositol 3,4,5-triphosphate (PIP_3_) production^20^. Vinculin is present at FAs in a force-dependent manner and increases cell stiffness^21^. To evaluate the maturation of FA-actin complex, we compared the vinculin expression and monitored PIP_3_ levels using a GFP-tagged PIP_3_ reporter (GFP-Grp1-PH) in static and stretch conditions in CTRL and *Thbs1*KO cells. PIP_3_ was enriched on stress fibers in stretched-CTRL cells (Online Fig. VI) and vinculin was deposited onto the tip of actin stress fibers which anchored FAs in stretched-CTRL cells, whereas stretched*-Thbs1KO* cells did not show recruitment of vinculin onto FA-actin complex (Fig. 3E). To confirm this observation, we assessed mechanical properties of *Thbs1KO* cells using atomic force microscopy (AFM) (Fig. 3F). Since cell tension positively correlates with adhesion size and its stiffness^22^, we analyzed morphological changes and stiffness on the stress fibers. Stretched-*Thbs1KO* cells showed a significant decrease in stiffness (Young’s modulus) on the stress fibers compared with stretched-CTRL cells, although height of stretched-*Thbs1KO* cells was comparable to that of stretched-CTRL cells (Fig. 3G). Taken together, our findings suggest that Thbs1 is required for maturation of FA-actin complex and maintenance of cellular tension thereby correctly orienting actin fibers in response to cyclic stretch.

### Deletion of thrombospondin-1 alters stretch-induced YAP-target gene expression

To understand the molecular basis of Thbs1 contribution in mechanotransduction, we performed RNA sequencing (RNA-seq) using CTRL and *Thbs1KO* cells in static and cyclic stretch conditions. Venn diagrams display the overlapping genes in each condition (Fig. 4A). In static condition, RNA-seq revealed 38 genes and 47 genes that were differentially regulated in CTRL and *Thbs1KO* cells, respectively. Conversely, 126 genes in CTRL cells and 163 genes in *Thbs1KO* cells were differentially expressed under cyclic stretch. PCA analysis identified 4 distinct groups according to conditions and genotypes (Fig. 4B). To evaluate stretch-induced gene expression between CTRL and *Thbs1KO* cells, we focused on the 126 genes (9.2%) in CTRL and 163 genes (11.9%) in *Thbs1KO* cells (Figs. 4C, D) that were differentially expressed under cyclic stretch. Among these genes, GO enrichment analysis revealed the abundant expressions of genes involved in response to stimulus (Online Fig. VII-A). We therefore used known gene groups involved in mechanical stress-induced responses, including YAP^23^, Notch^24, 25^, hypoxia-inducible factor-1a (HIF1A)^26^, ER stress response^27^, and oxidative stress response^28, 29^ as keywords to further characterize 289 genes via IPA combined with RNA-seq analysis (Online Fig. VII-B). With the exception of gene encoding Mesothelin (*Msln*), the expressions of YAP target genes were markedly downregulated in *Thbs1KO* cells, suggesting that Thbs1 strongly controls YAP target genes in stretch condition (Figs. 4E-I). Collectively, our data indicated that deletion of *Thbs1* affected stretch-induced gene expression, particularly YAP target genes, implying that Thbs1 may be a critical component of the YAP signaling pathway.

**Figure 4.**
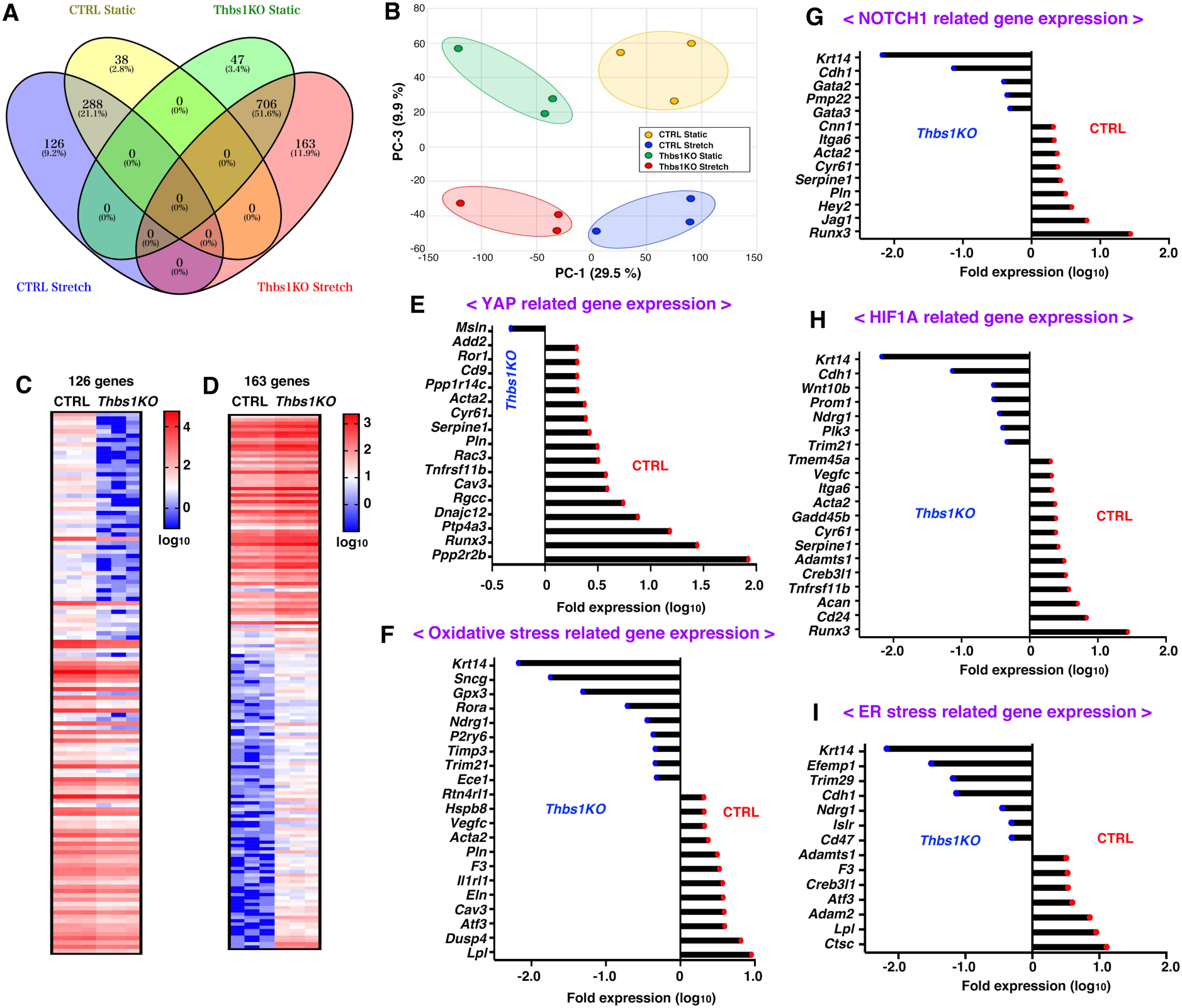
RNA expression profiles between CTRL and *Thbs1KO* cells under cyclic stretch. RNA-seq of CTRL and *Thbs1 KO* cells under static or cyclic stretch. **A**, Venn diagram summarizing the overlap genes among indicated groups. **B**. PCA plots showing distinct groups according to the condition and genotype. PC-1 (x-axis, 29.5%) and PC-3 (y-axis, 9.9%). **C-D**, Heat map showing the individual log_10_ values of gene expression from 3 independent experiments in stretched-CTRL cells (126 genes in A) in comparison to stretched-*Thbs1KO* cells (C), and those of stretched-*Thbs1KO* cells (163 genes in A) in comparison to stretched-CTRL cells (D). **E-I**, Among 289 genes in C-D, fold expression changes (log_10_) are shown as CTRL (red)/*Thbs1KO* (blue) in YAP-related genes (E), oxidative stress-related genes (F), Notch1-related genes (G), Hif1A-related genes (H), and ER stress-related genes (I).

### Thrombospondin-1 regulates nuclear shuttling of YAP via integrin αvβ1 in a Rap2-dependent manner

To elucidate the role for Thbs1 in the mechanical regulation of YAP, we examined YAP localization with or without Thbs1 under cyclic stretch. Consistent with a previous report^23^, cyclic stretch induced nuclear shuttling of YAP in CTRL cells (Fig. 5A). Although YAP retained in cytoplasm after cyclic stretch in *Thbs1KO* cells (Fig. 5A), exogenously added recombinant human THBS1 (rhTHBS1) promoted nuclear shuttling of YAP after cyclic stretch (Fig. 5B and Online Fig. VIII). Since nuclear shuttling of YAP is regulated by a kinase cascade of the Hippo pathway, we examined p-YAP and p-LATS levels using Western blotting. In CTRL cells, p-YAP and p-LATS were decreased in stretch condition whereas p-YAP and p-LATS remained unchanged in *Thbs1KO* cells (Fig. 5C). These observations indicate that Thbs1 controls nuclear shuttling of YAP in a Hippo pathway-dependent manner. The Ras related GTPase, Rap2 has recently been identified as a negative regulator of the Hippo pathway and to control YAP activation induced by ECM stiffness^30^. To determine whether stretch- and Thbs1-induced nuclear shuttling of YAP is associated with Rap2 activity, we measured active Rap2 (Rap2-GTP) levels in CTRL or *Thbs1KO* cells in static and stretch conditions using Ral-GDS-RBD pull down assays. As shown in Fig. 5D, active Rap2 decreased in CTRL cells after stretch but not in *Thbs1KO* cells. To further determine the relationship between Rap2 activity and nuclear shuttling of YAP, CTRL or *Thbs1KO* cells were transfected with myc-Rap2A (WT), myc-Rap2A (G12V) dominant active mutant, or myc-Rap2A (S17N) dominant negative mutant. Forty-eight hours after transfection, cells were subjected to cyclic stretch (1.0 Hz, 20% strain) for 8 h and immunostained with Myc, YAP, and DAPI. Rap2 (WT) overexpression showed no effects on the localization of YAP in CTRL or *Thbs1KO* cells (Fig. 5E). Rap2A activation (G12V) cancelled nuclear shuttling of YAP under cyclic stretch in CTRL cells. In contrast, Rap2A inactivation (S17N) forced YAP nuclear shuttling with or without Thbs1, overriding the regulation of Thbs1. These data suggested that Thbs1 controls nuclear shuttling of YAP in a Rap2A-dependent manner.

**Figure 5.**
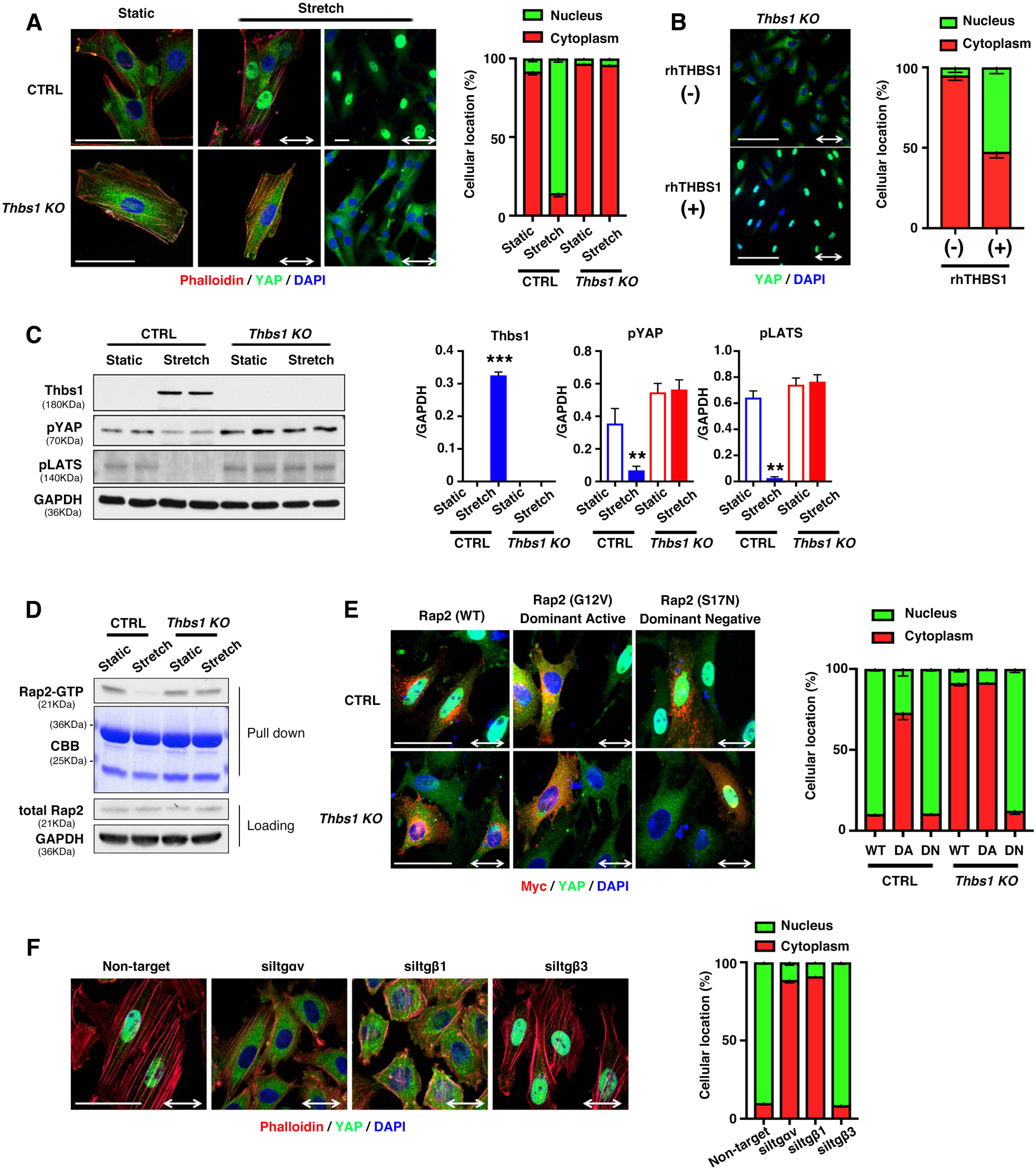
Thbs1 mediates nuclear shuttling of YAP via integrin αvβ1 in a Rap2-dependent manner. **A**, Representative immunostaining of CTRL or *Thbs1KO* cells in static or stretch condition with YAP (green). Phalloidin (red) and DAPI (blue). Two-way arrows indicate stretch direction. Bars are 50 µm. Quantification of YAP localization shown in right. 150-200 cells were evaluated in each experiment (n=3). YAP localizes in nuclear (green) or cytoplasm (red). **B**, *Thbs1KO* cells treated with (+) or without (-) recombinant human THBS1 (rhTHBS1;1µg/ml) following 24 h of serum starvation, then subjected to uniaxial cyclic stretch (20% strain; 1Hz) for 8h. Representative immunostaining with YAP (green) and DAPI (blue). Bars are 50 µm. Quantification of YAP localization shown in right. 80-120 cells were evaluated in each experiment (n=3). **C**, Western blot shows Thbs1, pYAP and pLATS levels in CTRL and *Thbs1KO* cells in static or stretch condition (n=3). Quantification graphs shown in right. Bars are means ± SEM. \*\**P*<0.01, \*\*\**P*<0.001, one-way ANOVA. **D**, Pull down of GTP-bound Rap2 from CTRL and *Thbs1KO* cells in static and stretch condition using Ral-GDS-RBD agarose beads (n=2). **E**, Overexpression of Myc-Rap2A(WT), Myc-Rap2A(G12V); dominant active or Myc-Rap2A(S17N); dominant negative in CTRL and *Thbs1KO* cells in static or stretch condition (n=3). Representative immunostaining with Myc (red), YAP (green), and DAPI (blue). Two-way arrows indicate stretch direction. Bars are 50 µm. Quantification of YAP localization shown in right. 50-100 cells were evaluated in each experiment (n=3). YAP localizes in nuclear (green) or cytoplasm (red). **F**, Scramble or targeted siRNA-treated rat vascular SMCs were subjected to cyclic stretch (n=3). Representative immunostaining with YAP (green), Phalloidin (red), and DAPI (blue). Two-way arrows indicate stretch direction. Bars are 50 µm. Quantification of YAP localization shown in right. 150-200 cells were evaluated in each experiment (n=3). Quantitative polymerase chain (qPCR) data for confirming the knockdown of *Itgαv, Itgβ1* and *Itgβ3* are provided in Online Fig. X.

Tension on actin fibers is also important for regulation of YAP, dependent or independent of the formation of FAs^31-33^. We assessed how actin dynamics and inhibition of FAK affects stretch-induced nuclear shuttling of YAP. CTRL cells were subjected to cyclic stretch for 8 h with the actin polymerization inhibitor, cytochalasin D (CytoD) or actin depolymerization inhibitor, jasplakinolide (Jasp) or FAK inhibitor PF-00562271 (FAKi). The efficiency of the treatment was evaluated by measuring the orientation angle (θ) of the long axis of the ellipse relative to the cyclic stretch direction (Online Fig. IX-A, B). Although CytoD or Jasp showed no effects on nuclear shuttling of YAP under cyclic stretch, YAP remained in the cytoplasm in FAKi-treated cells (Online Fig. IX-C, E). Furthermore, overexpression of slingshot1 (Ssh1, Ssh1OE), a phosphatase of cofilin, induced depolymerization of actin fibers through dephosphorylation of cofilin; however, YAP localization was not affected in Ssh1OE cells under cyclic stretch (Online Fig. IX-D, E). These observations suggest that stretch-induced nuclear shuttling of YAP is dependent on FAs. To confirm this, we performed small interfering RNA (siRNA)-mediated knockdown (KD) of integrin αv (*siItgαv*), β1 (*siItgβ1*) and β3 (*siItgβ3*) along with scramble siRNA (Online Fig. X) followed by cyclic stretch. As expected, knockdown of integrin αv or β1 was sufficient to prohibit stretch-induced nuclear shuttling of YAP (Fig. 5F). We next investigated whether matrix proteins in general serve as an extracellular regulator of YAP under cyclic stretch. Since fibronectin has been shown to regulate the Hippo pathway through FAK-Src-PI3K during cell spreading^34^ and a ligand for integrin αvβ1, we knocked down fibronectin in SMCs and subjected them to cyclic stretch. As Online Fig. XI shows, fibronectin knockdown had no effect on nuclear shuttling of YAP in response to stretch. We also assessed whether Thbs1-mediated nuclear shuttling of YAP is observed in endothelial cells. Using human umbilical vein endothelial cells (HUVECs), cyclic stretch did not induce phosphorylation of Paxillin or localization of Thbs1 to FAs, even though Thbs1 was secreted into CM (Online Figs. XII-A, B). Actin fibers aligned in a perpendicular position and contact inhibition was reduced by cyclic stretch, but YAP remained in the cytoplasm of HUVECs (Online Figs. XII-C, D), suggesting that regulation of YAP by Thbs1 under cyclic stretch condition is specific to vascular SMCs. Taken together, these data indicate that stretch-induced, Thbs1-mediated nuclear shuttling of YAP is dependent on integrin αvβ1 and independent of altered actin fibers.

### Thrombospondin-1/integrin/YAP signaling is involved in mechanical stress-induced vascular pathology

An in vivo model was used to evaluate whether Thbs1-mediated YAP regulation is involved in the pathogenesis of vascular diseases. We previously reported that transverse aortic constriction (TAC) induced Thbs1 expression^12^. Concordantly, Thbs1 was induced in ECs, SMCs and adventitial cells after TAC was performed in wild-type (WT) mice and proximity ligation assay (PLA) showed the interaction between Thbs1 and integrin β1 in these cells (Fig. 6A). We next evaluated the effect of loss of Thbs1/integrin β1 in TAC-induced aortic remodeling using *Thbs1* null (*Thbs1KO*) mice. Whereas 100% of WT mice survived at 5 weeks after TAC (n=13), 31.9% (7/22) of *Thbs1KO* mice died, in which 3 showed aortic rupture (Fig. 6B and Online Fig. XIII). The heart weight and body weight ratio (HW/BW) was significantly higher in TAC-operated *Thbs1KO* mice than that of WT mice in accordance with the previous report^35^ (Fig. 6C). In addition, TAC-operated *Thbs1KO* aortas had enlarged aortic lumen (Fig. 6D) and dissected aortas showed severely disrupted elastic fibers (Fig. 6E). Morphometric analysis showed a significant increase in the internal elastic lamella (IEL) perimeter, outer perimeter and total vessel area in TAC-operated *Thbs1KO* aortas compared to TAC-operated WT aortas (Fig. 6F). Immunostaining showed strong expression of YAP and its target connective tissue growth factor (CTGF) in TAC-operated WT aortas, especially medial layers and adventitia (Figs. 6G, H). However, their expression was hardly observed in the medial layers of TAC-operated *Thbs1KO* aorta, although the expression in the adventitia was evident in *Thbs1KO* aortas. These data suggest the possibility that Thbs1/integrin/YAP signaling in SMCs plays an important role in adaptive remodeling of the aortic wall in response to pressure overload.

**Figure 6.**
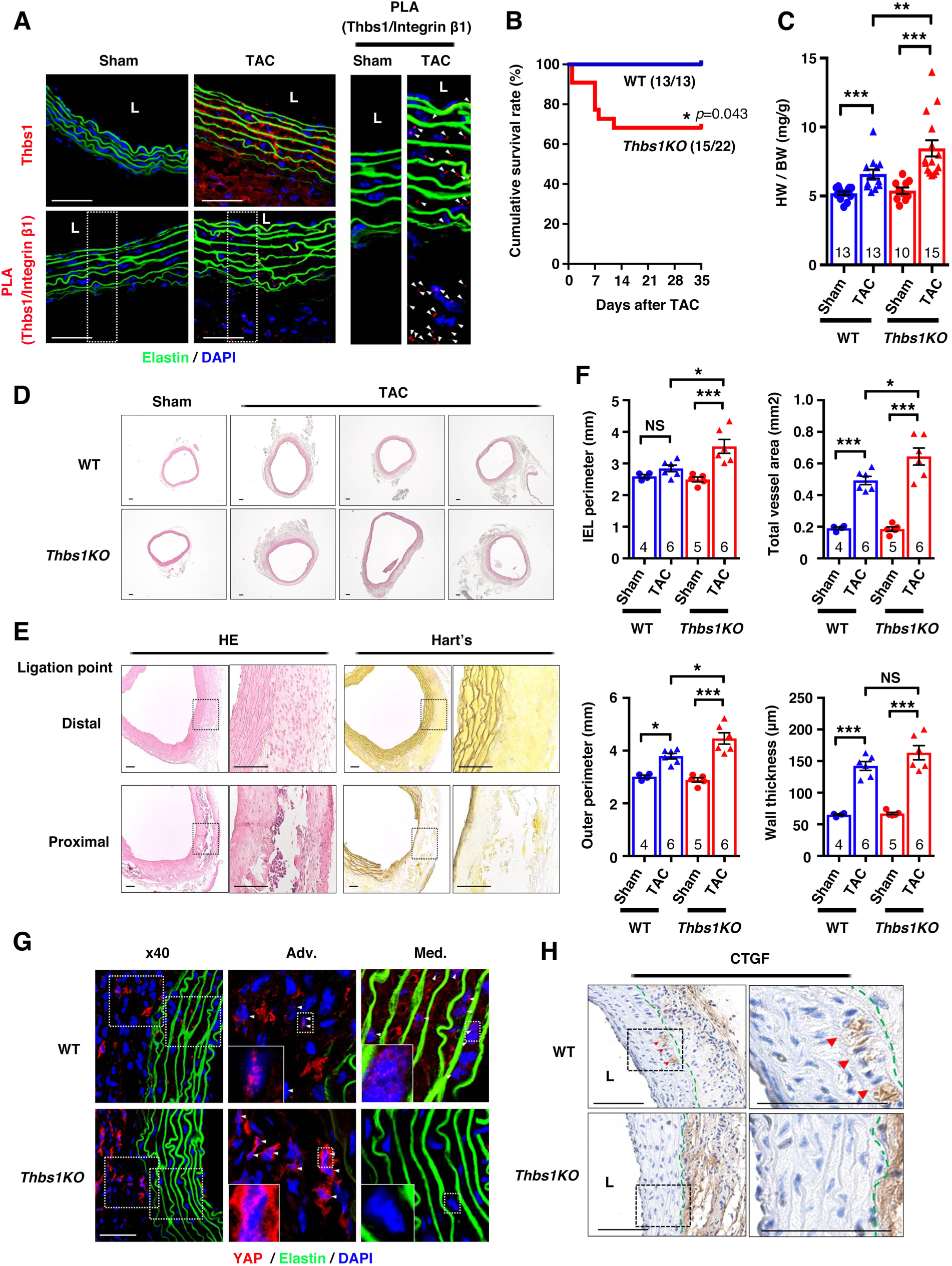
Thbs1 deficiency in mice results in aortic dissection under pressure overload. **A**, Immunostaining of Thbs1 (red) in sham or TAC-operated aortas in WT mice (upper). PLA showing the interaction between Thbs1 and Integrin β1(red dots in bottom). Right panels show high-magnified images of white-dashed boxes in the bottom panels. Arrowheads (white) show PLA cluster. DAPI (blue) and auto-fluorescence of elastin (green). n=3. Bars are 50 µm. **B**, Kaplan-Meier survival curve for WT (n=13) and *Thbs1KO* mice (n=22) after TAC. \**P*=0.043, Gehan-Breslow-Wilcoxon test. **C**, Heart weight to body weight ratio (HW/BW) in sham or TAC-operated mice. Bars are mean ± SEM. \*\**P*<0.01, \*\*\**P*<0.001, 1-way ANOVA. Number of animals is indicated in each bar. **D**, H&E staining of the ascending aortas for histological comparison between sham and TAC-operated aortas. Bars are 100 µm. **E**, Aortic dissection was observed in *Thbs1KO* ascending aorta proximal to the constriction. Bars are 100 µm. **F**, Morphometric analysis showing IEL perimeter, outer perimeter, total vessel area and wall thickness. Bars are mean ± SEM. \**P*<0.05, \*\**P*<0.01, \*\*\**P*<0.001, 1-way ANOVA. Number of animals is indicated in each bar. **G**, Immunostaining of TAC-operated aortas with YAP (red) in WT and *Thbs1KO* mice. High-magnified images of white-dashed boxes in the adventitia (Adv.) and medial (Med.) layers in left are shown. DAPI (blue) and auto-fluorescence of elastin (green). n=3. Bars are 50 µm. **H.** Representative images of immunohistochemistry for connective tissue growth factor (CTGF) in TAC-operated aortas of WT and *Thbs1KO* mice (n=3 per genotype). Outer elastic lamella as a border between the medial layer and adventitia shown in green-dotted line. Arrowheads (red) indicated CTGF positive cells in medial layer. Bars are 100 µm.

Finally, we employed carotid artery ligation injury to address the contribution of Thbs1/integrin/YAP signaling in flow secession-induced neointima formation (Fig. 7A). Thbs1 has been shown to be required for neointima formation^36^ and YAP mediates phenotypic modulation of SMCs during neointima formation^37^. Time course analyses of the ligated arteries revealed that Thbs1 was upregulated in the inner layer at 1 week after ligation when YAP expression was not evident (Fig. 7B and Online Fig. XIV). At 2 weeks after ligation, nuclear YAP was peaked in the developing neointima (Fig. 7C) and co-expressed with Thbs1 (Fig. 7B and Online Fig. XV). At 3 weeks, Thbs1 and YAP expression decreased as the neointima became stabilized (Fig. 7B and Online Fig. XVI). CTGF, a YAP target, was highly expressed in the neointima (Online Fig. XVII). PLA showed the interaction between Thbs1 and integrin β1 in the neointima (LCA in Fig. 7D), but not in the contralateral artery (RCA in Fig. 7D). To confirm the relationship between Thbs1/integrin/YAP and neointima formation, we injected the lentivirus (LV)-encoding *Thbs1* siRNA (siThbs1) into WT mice through tail vein injection at 2 weeks after ligation and evaluated neointima at 3 weeks. GFP-LV injection from the tail vein showed a clear expression of GFP in the neointima, endothelial cells of the right carotid artery, and the liver (Online Fig. XVIII), confirming the delivery of siRNA into the neointima. As we expected, the neointima was not observed in siThbs1 virus-treated animals, similar to *Thbs1KO* mice, and YAP levels and nuclear localization were markedly decreased as Thbs1 expression decreased (Figs. 7C, E).

**Figure 7.**
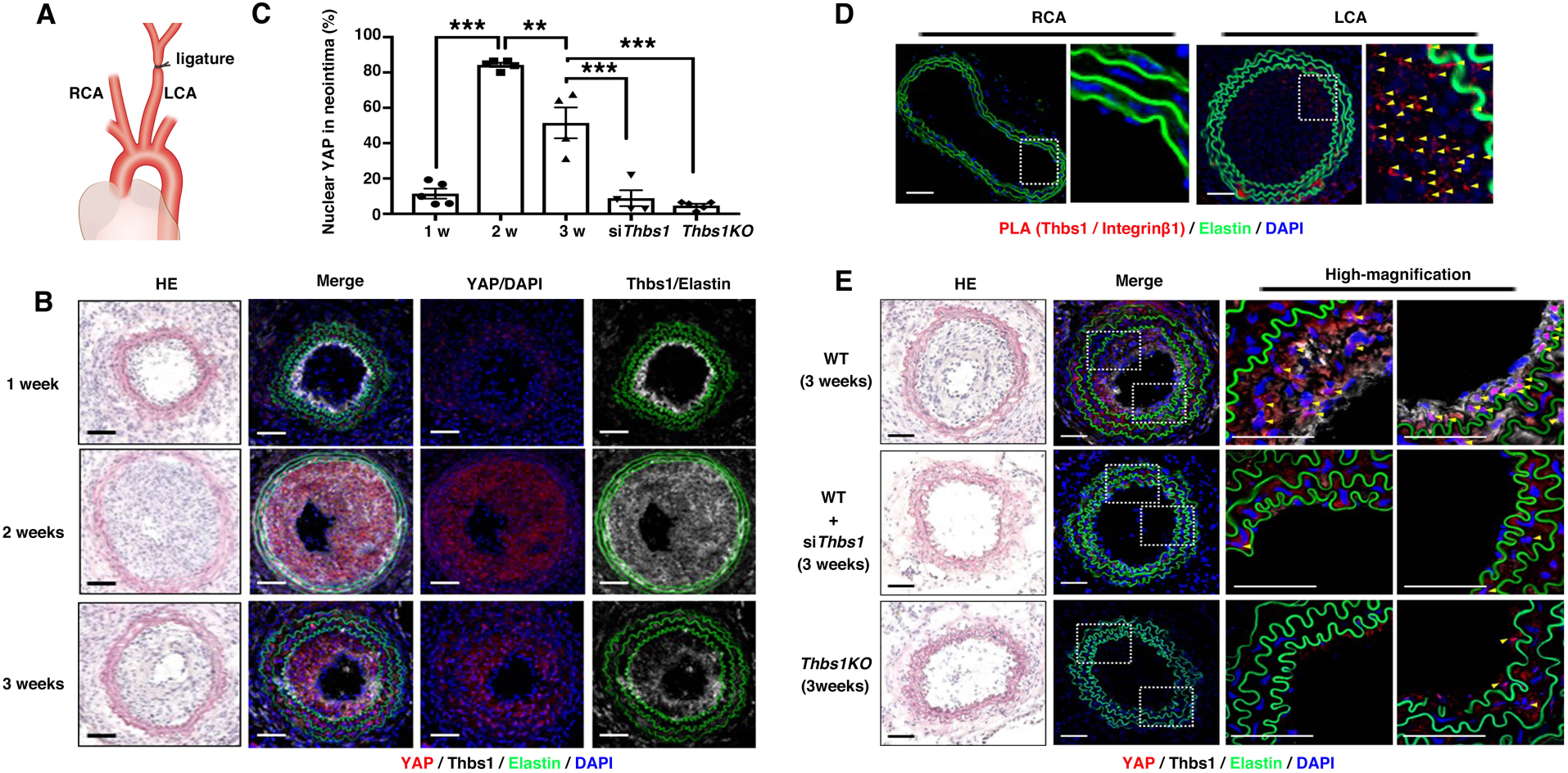
Thbs1 regulates YAP expression during neointima formation upon carotid artery ligation. **A**, Diagram of left carotid artery (LCA) ligation. RCA: right carotid artery. **B**, Immunostaining of cross sections of ligated arteries with YAP (red) and Thbs1 (white) at 1 week (n=5), 2 weeks (n=5) and 3 weeks (n=4) after ligation in WT mice. DAPI (blue) and auto-fluorescence of elastin (green). Bars are 50 µm. **C**, Quantification of co-localization with YAP and DAPI using Imaris colocalization software. Bars are mean ± SEM. \**P* < 0.05, \*\**P* < 0.01, \*\*\**P* < 0.001, one-way ANOVA. **D**, PLA showing cluster of Thbs1/Integrin β1(red dots, yellow arrowheads) at 3 weeks after ligation in RCA and LCA. DAPI (blue) and auto-fluorescence of elastin (green). Bars are 50 µm. n=3. **E**, Cross sections of ligated arteries at 3 weeks post-injury in WT (n=4), si*Thbs1*-treated WT mice (n=4), and *Thbs1KO* (n=5) mice. Immunostaining with YAP (red), Thbs1 (white). DAPI (blue) and auto-fluorescence of elastin (green). Note neointima formation in WT arteries. Yellow arrowheads show nuclear localization of YAP. Bars are 50 µm.

## Discussion

In this study, we identified Thbs1 as a critical molecule involved in mechanotransduction and extracellular regulation of YAP, leading to dynamic remodeling of the blood vessels. We found that Thbs1, once secreted in response to mechanical stress, binds to integrin αvβ1 and aids in the maturation of FA-actin complex, downregulating Rap2 activity, thereby mediating activation of YAP. Loss of Thbs1 decreases cell stiffness in vitro and deletion or downregulation of Thbs1 in mice alters cellular behavior in response to mechanical stress in a context dependent-manner. Taken together, our study demonstrates that Thbs1/integrin/YAP signaling pathway plays a critical role in vascular remodeling in vivo (Fig. 8).

**Figure 8.**
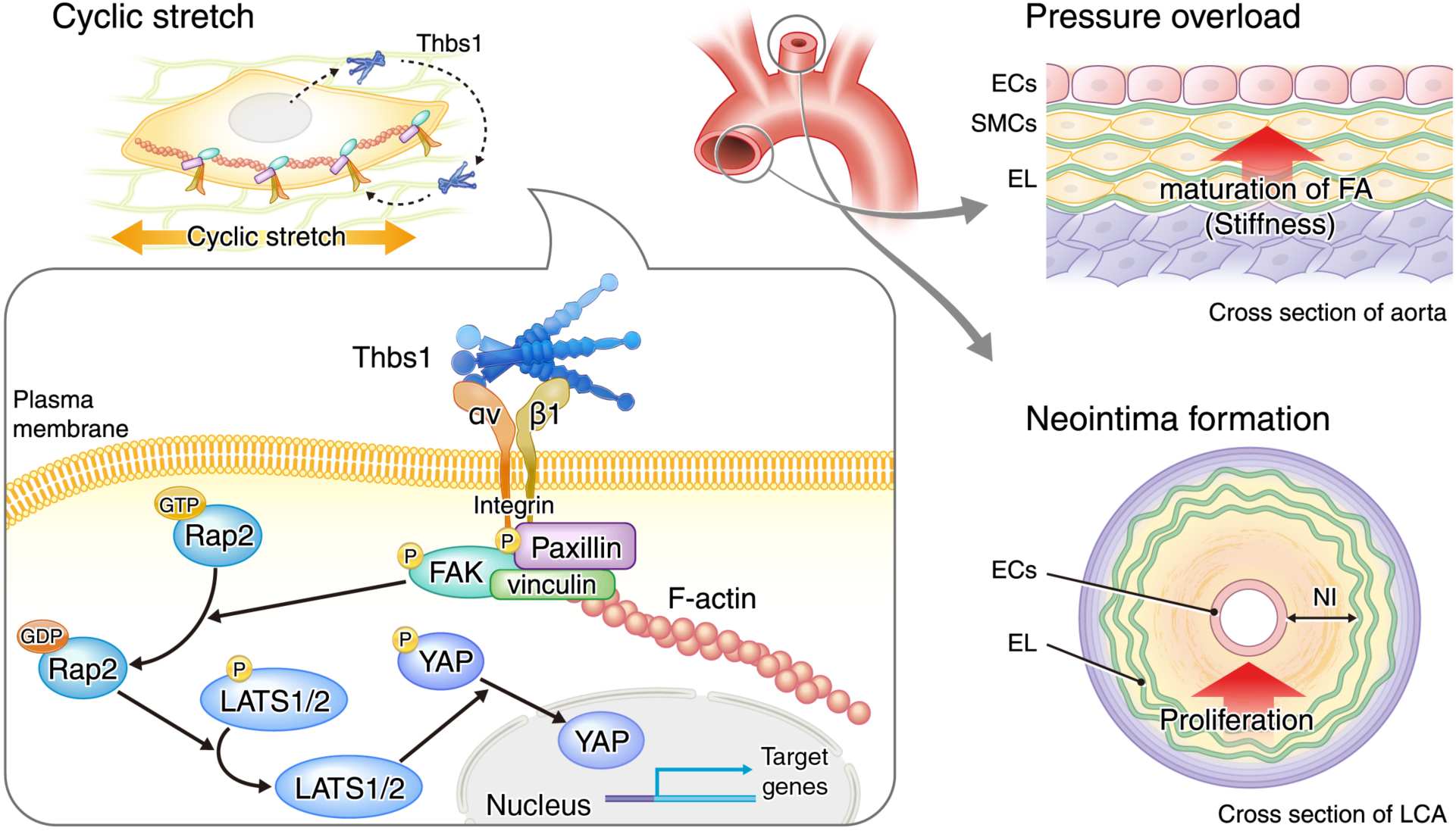
A model illustrating the Thbs1/integrin/YAP signaling pathway in remodeling of vessel walls. Cyclic stretch induces secretion of Thbs1, which binds to integrin αvβ1 and aids in the maturation of FA-actin complex, thereby mediating nuclear shuttling of YAP via inactivation of Rap2 and Hippo pathway. In vivo, Thbs1/integrins/YAP signaling leads to maturation of FA and an increase in stiffness of SMCs after TAC-induced pressure overload to protect the aortic wall. In contrast, Thbs1/integrins/YAP signaling leads to neointima (NI) formation upon flow cessation by carotid artery ligation. ECs: endothelial cells, SMCs: smooth muscle cells, EL: elastic lamella, LCA: left carotid artery.

### Mechanical stress and micro- and macro-remodeling of the vessel wall by ECM

Vascular ECM, a major component of the vessel wall, is subjected to repeated mechanical stress throughout life. Due to its long half-life (for example, 50-70 years for elastic fibers of human artery^38^), the vascular ECM is considered to be difficult to regenerate once it is damaged by physical and/or biochemical insults. Although ECM synthesis via mechanical stretch has been previously described^39, 40^, its biological significance, whether it contributes to micro- or dynamic remodeling of the vessel wall, and how it is connected to the mechanotransduction cascade are not completely understood. Using mass spectrometry and GO analysis, we found that cyclic stretch induced secretion of various ECM proteins involved in cell adhesions, elastic fiber formation and TGFβ signaling. Using IPA, Thbs1 was identified as one of the key molecules involved in these biological processes. Interestingly, mechanical stress and secretary pathway are interconnected since BFA-treatment or deletion of *Thbs1* led to the alteration of mechano-response in SMCs and disrupted alignment of cells in response to cyclic stretch. Our observations indicate that mechanical stress induces secretion of Thbs1, which initiates mechanotransduction and activates integrin/YAP pathway, thereby remodeling the vessel wall to accommodate the changes in mechanical microenvironment. Thbs1 and integrins are essential components of matrix mechanotransduction, serving as biomechanical ligand and sensors of the microenvironment.

### Thbs1 as an extracellular mediator of YAP signaling pathway

We showed that mechanical stretch-induced, secreted Thbs1 directly bound to integrin αvβ1 and co-localized with p-Paxillin, and facilitated maturation of FA complex by depositing vinculin at the tip of actin fibers. Currently, the precise mechanism of how secreted-Thbs1 is recruited to integrin αvβ1 is unclear. However, we speculate that Thbs1 ligation to integrins activates downstream molecules and lead to maturation of FAs and strengthen actin fibers. Interestingly, a loss of Thbs1 decreased cell stiffness and altered YAP-target gene expression after cyclic stretch. We further showed that stretch downregulated Rap2 activity in CTRL cells, whereas Ras2 activity remained high in the absence of Thbs1, which forced YAP in the cytoplasm. Our observation agrees with the recent study by Meng et al demonstrating that Rap2 negatively regulates nuclear shuttling of YAP in a Hippo pathway dependent manner. In their study, the authors proposed a cascade of signaling triggered by FAs → PLCγ1 (phospholipase C gamma 1) → PtdIns(4,5)P_2_ (phosphatidylinositol 4, 5-bisphosphate) → PA (phosphatidic acid) → PDZGEF (RAPGEF) → Rap2 → MAP4K → LATS → YAP^30^, and increased matrix stiffness led to inactivation of Rap2 and nuclear shuttling of YAP. It was also proposed that this signaling axis works in parallel with RhoA, which mediates cytoskeletal tension during cell spreading ^5, 32^. Our data showed that stretch-induced YAP activation is Rap2-dependent and is not influenced by altered actin fibers. Therefore, it is possible that Thbs1 mediates nuclear shuttling of YAP by inactivating Rap2 through altering the composition of FA or by controlling FA assembly ^41^. It has previously been described that a mutant cell line lacking YAP showed decreased Thbs1 levels along with downregulation of YAP-target genes and FA molecules, indicating that Thbs1 may act as a feedforward driver for YAP signaling^41^. Interestingly, the proteolytic remodeling of laminin-1 has been shown to modulate cancer cell awakening through integrin/FAK/ERK/MLCK (myosin light-chain kinase)/YAP signaling^42^, further supporting the idea that an ECM can regulate YAP signaling in various biological conditions.

### Role of matrix mechanotransduction in vessel wall remodeling

Recent studies have shown that ECM mechanical cues mediate cell proliferation, differentiation, autophagy and lipid metabolisms^1, 43-45^. Thbs1 is present at low levels in the postnatal vessels and upregulated in various vascular diseases such as atherosclerosis^46^, ischemia reperfusion injury^47^, aortic aneurysm^13^ and pulmonary arterial hypertension^48^. YAP is also known to play a role in an emergency stress response to maintain tissue homeostasis^49, 50^. As we describe here, suppression of Thbs1/integrin/YAP signaling using *Thbs1KO* mice showed maladaptive remodeling in response to pressure overload. In WT mice, YAP was induced and activated in adventitial cells and SMCs, leading to proliferation and hypertrophy, respectively, and maintenance of wall tension during aortic wall dilatation. In contrast, *Thbs1KO* mice showed high mortality after TAC and caused aortic dissection without upregulation of YAP most likely in SMCs. These data suggest that formation of FAs by Thbs1 and induction of Thbs1/integrin/YAP signaling plays a protective role in response to pressure overload by presumably increasing cell stiffness. On the other hand, Thbs1/integrin/YAP signaling mediates proliferation of neointimal cells upon carotid artery ligation, exhibiting the adverse effect on vascular remodeling. It is of note that inhibition of Thbs1 by lentivirus-encoding siRNA was sufficient to suppress neointima formation. These observations emphasize the highly context-dependent action of Thbs1/integrins/YAP in vessel wall remodeling. It is plausible that different mechanical stress induces a distinct subset of YAP target genes and leads to vascular remodeling.

We have recently reported that Thbs1 is highly upregulated in the aneurysm lesions in *Fbln4* mutant mice and patients with thoracic aortic aneurysms (TAA)^13^, and deletion of *Thbs1* or administration of phosphoinositol-3-kinase (PI3K)-inhibitor prevented the aneurysm formation ^12, 13^. Since PI3K is also known to link YAP signaling pathway^51, 52^, it is possible that suppression of *Thbs1* or PI3K may alter YAP levels in *Fbln4* mutant mice and affected YAP target genes. Another possibility is that the effect of Thbs1 in *Fbln4* mutant aortic aneurysms is mechanistically different from that of physiological remodeling, where mutant SMCs cannot properly sense mechanical stress due to disrupted elastic lamina-SMC connections^13^. In this case, upregulation of Thbs1 cannot induce nuclear shuttling of YAP due to inability to form FAs, thereby the effect of Thbs1 may be independent of YAP signaling even in the presence of high Thbs1. It is critical, however, to consider whether inhibition of Thbs1 affects Thbs1/integrins/YAP pathway and potentially alters the remodeling processes in the vessel wall.

## Nonstandard Abbreviations and Acronyms

ECM: Extracellular matrix
p: Phosphorylated
SMC: Smooth muscle cell
EC: Endothelial cell
FAs: Focal adhesions
YAP: Yes-associated protein
Thbs1: Thrombospondin-1
FAK: Focal adhesion kinase
BFA: Brefeldin A
IPA: Ingenuity Pathway Analysis
PLA: Proximity Ligation Assay
AFM: Atomic force microscopy
RNA-seq: RNA sequencing
DAPI: 4’, 6-diamidino-2-phenylindole
CM: Conditioned media
TAC: Transverse aortic constriction
IEL: Internal elastic lamella
LCA: Left carotid artery
CTGF: Connective tissue growth factor
GAPDH: Glyceraldehyde-3-phosphate dehydrogenase

## Acknowledgement

We thank S. Sato, K. Sugiyama and HT. Hang for technical assistance. We acknowledge technical support from JD. Kim, M. Muratani and Tsukuba i-Laboratory LLP for RNA-seq analysis. We appreciate Z.-P. Liu, K. Ohashi, and T. Mayers for critical reading of the manuscript. We thank Mayumi Mori for assistance with graphics.

