## Supplemental information for "Matrix mechanotransduction mediated by thrombospondin-1/integrin/YAP signaling pathway in the remodeling of blood vessels"

### Detailed Methods

***Cell culture and reagents.*** Rat vascular SMCs (Lonza, R-ASM-580, isolated from the aorta of 150-200 grams adult male Sprague-Dawley rats), were grown in DMEM with 20% (v/v) fetal bovine serum and 1 x Antibiotic-Antimycotic (Thermo Fisher Scientific). HUVECs were grown in HuMedia-MvG (KURABO) supplemented with 5% (v/v) fetal bovine serum and growth factors. The cell lines were used between passage 7 and 12 and tested for mycoplasma contamination using Mycoplasma detection set (TaKaRa, #6601) following the manufacture's protocol (result provided in Online Fig. XIX). Brefeldin A (B5936) was purchased from Sigma-Aldrich. PF-00562271, FAK inhibitor (S2672) was purchased from Selleck. Cytochalasin D (037-17561) and Latrunculin A (125-04363) were purchased from Wako. Jasplakinolide (AG-CN2-0037-C050) was purchased from AdipoGen Life Sciences. Recombinant human thrombospondin-1 was purchased from R&D Systems (3074-TH-050).

***Plasmid.*** GFP-Grp1-PH plasmids were provided by T. Ijyuin (Kobe University, Japan)[1]. Myc-tagged plasmids encoding mouse Rap2A(WT), Rap2A(G12V), and Rap2A(S17N) were kindly donated by K. Kariya (Ryukyu University, Japan)[2]. YFP-Ssh1 was donated by K. Mizuno (Tohoku university, Japan)[3].

***In vitro mechanical stretch.*** Cyclic stretch was performed using a uniaxial cell stretch system (Central Workshop Tsukuba University) following 24 h of serum starvation as described previously[4]. Rat vascular SMCs or HUVECs were plated (3 × $10^4$ cells) on silicon elastomer bottomed culture plates (SC4Ha, Menicon Life Science) coated with cell attachment factor, containing gelatin (S006100, Thermo Fisher Scientific) and subjected to cyclic stretch with a frequency of 1.0 Hz (60 cycles/min) and 20% strain for the indicated time.

***siRNA Transfection.*** Rat vascular SMCs were transiently transfected with 10 pmol for a final concentration of scramble (non-target) siRNA (Invitrogen, #12935110), *Itgav* siRNA (RSS310692), *Itgb1* siRNA (RSS302262), *Itgb3* siRNA (RSS308076) and *Fn1* siRNA (RSS303981) using RNAiMAX transfection reagent (Thermo Fisher Scientific) according to the manufacturer's instruction. All siRNAs were predesigned and purchased from Thermo Fisher Scientific. Cyclic stretch was performed 2 days after transfection.

***Western blot analysis.*** Rat vascular SMCs or HUVECs were dissolved in RIPA lysis buffer (Sigma) containing 1% protease inhibitor (Wako) and 1% phosphatase inhibitor (Wako). The lysates were mixed with 3 × SDS sample buffer with 2-mercaptoethanol (Wako) and boiled at 95 °C for 5 min, and then were subjected to SDS-PAGE. Proteins were transferred to a PVDF membrane (Millipore) and immunoblotted with antibodies indicated in Online Table I. Membranes were then incubated with respective anti-mouse or anti-rabbit HRP-conjugated secondary antibody (1:1000, Bio-Rad) and visualized with a chemoluminescence kit (Santa Cruz Biotechnology) or SuperSignal West Femto Maximum Sensitivity Substrate (Thermo Fisher Scientific).

***Quantitative Polymerase chain reaction (PCR).*** RNA was isolated from rat vascular SMCs using RNeasy Plus Micro Kit (QIAGEN) and 1 μg of total RNA was subjected to reverse transcription reactions using iScript™ Reverse Transcription Supermix (Bio-Rad). SYBR Green was used for Amplicon detection and gene expression was normalized to the expression of housekeeping genes Glyceraldehyde 3-phosphate dehydrogenase (GAPDH). PCR reactions were carried out in triplicate in a CFX96 real-time PCR detection system (Bio-Rad) with one cycle of 3 min at 95 °C, then 39 cycles of 10 sec at 95 °C and 30 sec at 55 °C. Levels of mRNA were determined using the ddCt method and expressed relative to the mean dCt of controls. Primer sequences are provided in Online Table II.

***Immunostaining.*** Rat SMCs or HUVECs, were fixed with 4% paraformaldehyde for 30 min, blocked in 5% BSA containing 0.1% Tween-20 for 1 h at 37 °C. Aortas and carotid artery were harvested and embedded in OCT (SAKURA Finetek USA Inc.) and snap frozen in liquid nitrogen. Cross sections of the mouse aortas or carotid arteries were immediately fixed with 4% paraformaldehyde for 30 min, blocked in 5% normal serum in which secondary antibody was raised, containing 0.1% Tween-20 for 1 h at 37 °C. The primary antibodies used are shown in Online Table I. Incubation was performed overnight at 4 °C. After washing, highly cross-absorbed Alexa 488, Alexa 546 or Alexa 647-conjugated secondary antibodies (Invitrogen) were added at a dilution of 1:200 for 2 h at 37 °C. Control experiments were performed by omitting the primary antibody. Slides were covered with Vectashield containing DAPI (Vector Laboratories) and viewed under a LSM 710 (ZEISS). Colocalization was measured on a single confocal section using the Colocalization-module of Imaris 9.2.1, 64-bit version (Bitplane AG).

***Proximity ligation assay (PLA).*** Duolink™ PLA (Sigma-Aldrich) was performed following the manufacturer's instruction. The primary antibodies used are shown in Online Table I. Alexa 488-phalloidin (Invitrogen) was added to visualize actin fibers. Incubations were performed overnight at 4 °C. After washing, the PLA probes; Duolink In Situ PLA probe Anti-Mouse PLUS (DUO92001) and Duolink In Situ PLA probe Anti-Goat MINUS (DUO92006) were added for 2 h at 37 °C. Duolink In Situ Detection Reagents Orange (Invitrogen) was used for detection. Slides were covered with Duolink In Situ Mounting Medium containing DAPI (DUO82040) and viewed under a LSM 710 (ZEISS).

***Immunoprecipitation.*** Immunoprecipitation was conducted using a Pierce Crosslink IP kit (Thermo Fisher Scientific). Antibodies against Thbs1 (A6.1 MS-421 and A4.1 MS-418, Neomarkers) or mouse normal IgG (10400C, Invitrogen) were crosslinked to protein A/G agarose beads and then incubated with protein lysates overnight at 4 °C. After washing and elution, input and IP products were analyzed using Western blotting.

***Rap2 activation assay.*** Rap2 activation assay was performed according to the manufacture's instruction (Cell Biolabs, STA-406-2) and previous studies[5, 6]. After cyclic stretch or static condition, CTRL and *Thbs1KO* rat vascular SMCs were lysed at 4 °C by Assay/lysis buffer containing 125 mM HEPES (pH7.5), 750 mM NaCl, 5% NP-40, 50 mM $MgCl_2$, 5 mM EDTA, 10% Glycerol and active Rap2 was pulled down from lysates using agarose beads conjugated to Ral-GDS-RBD and detected by Western blotting using mouse anti-Rap2 antibodies.

***Generation of Thbs1 knockout cell line.*** CRISPR-Cas9 plasmid, pSpCas9(BB)-2A-GFP (px458, #48138) was purchased from Addgene. The CRISPR design tool (http://crispr.mit.edu) was used to identify candidate sgRNA sequences of *Thbs1* gene. sgRNA sequence was PCR amplified and ligated into *Bbs*I site of px458. After transformation into *E.Coli,* the plasmid DNA was purified by using Midi prep kit (QIAGEN, Chatsworth, CA). Rat SMCs ($1 \times 10^5$ cells) were plated in a 24-well plate and transfected with px458_sgRNA plasmid. Forty-eight h after transfection, FACS (BD FACSAria™ II, BD) was performed to isolate GFP-positive cells. Following isolation, cells were plated in a 96-well plate by limiting dilution for a single-cell cloning. Gene mutations were confirmed using Sanger sequencing (3130xl Genetic Analyzer, Applied Biosystems).

***Secretome Analysis.*** Rat vascular SMCs were plated ($3 \times 10^4$ cells) on silicon elastomer bottomed culture plates (SC4Ha, Menicon Life Science) coated with cell attachment factor (Thermo Fisher Scientific). Following 24 h of serum starvation, cyclic stretch was performed (1.0 Hz, 20% strain) for 20 h with or without 1 μM BFA. Four mL of conditioned media (CM) was collected and concentrated to 300 μl using Amicon Ultra-3K (Millipore, UFC200324), from which 200 μl was enriched with acetone precipitation and the rest was used for Silver staining. The CM were diluted 4-fold with acetone and stored at -80 °C for 1 h followed by centrifugation at 20,400 × g for 20 min at 4 °C. After centrifugation, the supernatant was removed. Pellets were resolved in 50 μl of 100 mM Tris-HCl (pH 9.0) containing 12 mM sodium deoxycholate and 12 mM N-lauroylsarcosinate. The experiments were performed three times to prepare for proteomics (n=3).

***Sample preparation for proteomics*** The trypsin digestion was performed as described in a previously published study[7]. The samples were heated at 95 °C for 5 min and sonicated for 20 min, then reduced with 10 mM dithiothreitol at room temperature (RT) for 1 hour and alkylated with 55 mM iodoacetamide in the dark at RT for 1 h. The samples were diluted 5-fold with 50 mM ammonium bicarbonate and digested with lysyl endopeptidase at an enzyme/substrate ratio of 1:100 at RT for 3 h. Subsequently, the samples were digested with sequence-grade modified trypsin at an enzyme/substrate ratio of 1:100 at 37 °C for 16 h. Then, an equal volume of ethyl acetate was added to the sample and the mixture was acidified with trifluoroacetic acid (TFA) at a final concentration of 0.5%. The mixtures were centrifuged at 15,000 rpm for 2 min. The aqueous phases were dried up using a centrifugal evaporator (TOMY, Tokyo, Japan), and tryptic peptides were dissolved in 5% (v/v) acetonitrile with

0.1% TFA (v/v). The peptide samples were desalted with GL-Tip SDB and GL-Tip GC (GL Sciences). The eluents were dried up using a centrifugal evaporator and dissolved with 0.1% TFA for LC-MS/MS analysis.

***Nano LC-ESI-MS/MS analysis for global proteomics*** Samples were injected into Nano-LC system coupled with Triple TOF 5600 mass spectrometer (SCIEX, USA). Peptides were loaded onto 100 μm internal diameter x 20 mm length PepMap 100 precolumn (Thermo Fisher Scientific, USA) at 5 μl/min and separated by 75 μm internal diameter × 250 mm length PepMap RSLC column (Thermo Fisher Scientific, USA) packed with 2 μm C18 beads with 100 Å pores. The flow rate was 300 nL/min with a 107 min gradient of solvent B (100% acetonitrile and 0.1% formic acid) for separation on the reversed-phase column. The injected peptides were eluted with solvent A (0.1% formic acid in Milli-Q water) and solvent B. The gradient condition was set 2% B (0–3 min), changed to 2-25% B (3-63 min), increased to 25-50% B (63-78 min), maintained 98% B (80-85 min). Data acquisition was controlled using Analyst TF1.6 software (SCIEX, USA). For Information Dependent Acquisition (IDA), precursor ions were scanned from 300 to 1250, and product ions were scanned from 100 to 1600 with an accumulation time of 50 msec. The peptides were identified using ProteinPilot 4.5 (Sciex). For protein quantification, data was acquired with data independent acquisition mode (SWATH-MS) with valuable window for precursor ions. The acquired data was analyzed using PeakView Software version 2.1 (Sciex).

***Atomic Force Microscopy (AFM).*** AFM measurements were performed using a NanoWizard IV AFM (JPK Instruments-AG, Germany) mounted on top of an inverted optical microscope (IX73, Olympus, Japan) equipped with a digital CMOS camera (Zyla, Andor) as previously described[8]. The AFM quantitative imaging (QI) mode was then used to obtain a force-displacement curve at each pixel of 128 × 128 pixels (100 μm × 100 μm of measured area), using a precisely controlled high-speed indentation test using rectangular-shaped silicon nitride cantilevers with a cone probe (BioLever-mini, BL-AC40TS-C2, Olympus, Japan) at a spring constant of 0.08–0.10 N/m. The QI mode measurements were performed within 1 h after the transfer of the specimen cells to the AFM at room temperature RT (25 °C). These high-speed indentations were performed until reaching a pre-set force of 1 nN. Cell elasticity was calculated from the obtained force-displacement curves by applying a Hertzian model, approximating that the sample is isotropic and linearly elastic. Young's (elastic) modulus could be extracted by fitting all force-displacement curves with the following Hertzian model approximation:

$$F = \frac{2E \cdot \tan\alpha}{\pi(1-\nu^2)} \delta^2 \quad (1)$$

where $F$ is the applied force, $E$ is the elastic modulus, $\nu$ is the Poisson's ratio (0.5 for a non-compressible biological sample), $\alpha$ is the opening angle of the cone of the cantilever tip, and $\delta$ is the indentation depth of the sample. Using the results of the Hertzian model approximation, we identified the Z contact points (specimen surface) and the elastic modulus of the specimens at each pixel, and produced a surface topography map and elastic modulus map of the specimens to estimate cell height and mechanical tension of actin fibers.

***RNA-seq analysis.*** RNA was extracted from CTRL (n=3) and *Thbs1*KO (n=3) cells in static and stretch condition. Total RNA was isolated using TRIZOL reagent (Thermo Fisher) and quality of RNA was examined using an RNA 6000 Pico kit (Agilent) as previously described[9]. An amount of 500 ng total RNA was used for RNA-seq library preparation with NEB NEBNext Poly(A) mRNA Magnetic Isolation Module and ENBNext Ultra Directional RNA Library Prep Kit (New England Biolabs). Two × 36 base paired-end sequencing was performed with NextSeq500 (Illumina) by Tsukuba i-Laboratory LLP (Tsukuba, Japan). FASTQ files were analyzed using CLC Genomics Workbench (Version 10.1.1; Qiagen). Sequences were mapped to rat genome and quantified annotated genes. Transcript expression values were estimated as "reads per kilobase per million reads", and analyzed using Emperical Analysis of DGE tool (Qiagen).

***Bioinfomatics analysis.*** Functional network analyses of genes/proteins with statistically significant expression changes were performed using Ingenuity Pathway Analysis (Ingenuity Systems). Gene Ontology and Reactome pathway analyses were performed using GO Consortium (http://www.geneontology.org). Venn diagram was generated using Venny tool (http://bioinfogp.cnb.csic.es/tools/venny/) from BioinfoGP to identify the transcripts uniquely expressed in each condition.

***Mice.*** C57BL/6J (wild-type) mice were purchased from Charles River, Japan and *Thbs1* null mice were purchased from Jackson Laboratory (B6.129S2-$Thbs1^{tm1Hyn}$/J) and crossed with C57BL/6 mice for line maintenance. Male mice were used for experiments to exclude hormonal regulation by estrogen. All mice were kept on a 12 h/12 h light/dark cycle under specific pathogen free condition and all animal protocols were approved by the Institutional Animal Experiment Committee of the University of Tsukuba.

***Transverse aortic constriction (TAC).*** Eight-week-old male *Thbs1KO* mice or control littermates underwent TAC using a standard surgical protocol[10]. Briefly, anesthetized mice were placed in a spine position, and aortic constriction was achieved by tying a 6-0 silk suture (Natsume Seisakusho Co., Ltd.) against a 25-gauge blunt needle (TERUMO). For the sham group, the same operation was performed without ligating the aorta. Mice were kept for 5 weeks after TAC and the aortas were harvested.

***Carotid artery ligation.*** Eight-week-old male C57BL/6 or *Thbs1*KO mice were used for carotid artery ligation as previously described[11]. Briefly, the left common carotid artery was dissected free of connective tissues and permanently ligated proximally to the bifurcation. At 1, 2, or 3 weeks after surgery, mice were euthanized, perfused, and right and left carotid arteries were harvested and embedded in optimal cutting temperature compound. Serial cross sections were prepared from the ligature toward the aortic arch with 10 μm intervals until neointima disappeared.

***Production of lentivirus encoding siRNA and infection.*** Lentivirus expressing GFP (LV011-a) and siRNA targeting mouse *Thbs1* was purchased from Applied Biological Materials (abm) Inc. *Thbs1*-set siRNA (i043193) sequences were as follows, #si-RNA463 (AGCATCACGCTGTTTGTTCAAGAGGACCG); #siRNA1586 (GCAAAGACTGTGTTGG-CGATGTGACAGAA); #si-RNA2756 (AGAAGGACTCTGATGGTGATGGCCGAGGT); #siRNA3106 (TT-CTACGTTGTGATGTGGAAACAAGTCAC). siRNA lentivector and 3rd Generation Packaging System mix (LV053, abm Inc.) were co-transfected into 293T cells. Supernatant containing the lentiviral particles was collected 48 h after transfection and concentration of lentivirus was conducted using Lenti-X™ Concentrator. Transfection efficiency was calculated by using GFP-positive cells. For in vivo infection, the virus solution (100 μl, $1 \times 10^7$ IU/ml) was inoculated into the tail vein 2 weeks after carotid artery ligation.

***Immunohistochemistry, Histology and morphometric analysis.*** Aortas were harvested and fixed with 4% paraformaldehyde and embedded in paraffin. Five-micrometer sections were stained with H&E for routine histology and Hart's staining for detection of elastic fibers. For immunohistochemistry, primary antibody is shown in Online Table I. Incubation was performed overnight at 4 °C. After washing, anti-rabbit HRP-conjugated second antibody (1:1000, Bio-rad) was added for 1h. TaKaRa DAB Substrate (Takara) was used for detection and hematoxylin was used for counterstaining. Images were digitally captured with Axio Imager Z2 (ZEISS). Morphometric analysis was performed with NIH ImageJ software as previously described[4].

***Statistical analysis.*** All experiments are presented as means ± standard error of the mean (SEM). Statistical analysis was performed using Prism 7 (Graph Pad). $P < 0.05$ denotes statistical significance.

***Data availability.*** The acquisition of online accession numbers for the secretome data is jPOST ID : JPST000600/PXD013915 (http://jpostdb.org) and for the RNA-seq data is GSE131750 (GEO accession). All data supporting the findings of this study are available from the corresponding authors on reasonable request.

**Online Figures and Figure legends**

A

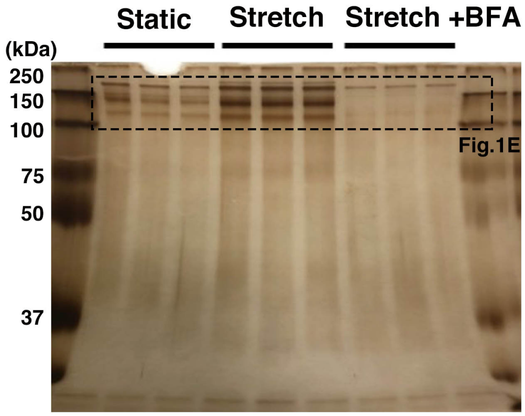


B

| | Proteins | Peptides |
|---|---|---|
| Static | 656 | 5344 |
| Stretch | 735 | 6831 |
| Stretch+BFA | 685 | 4912 |
| All | 870 | 8493 |

**Online Figure I. Secretome analysis of rat vascular SMCs under cyclic stretch. A,** Entire image of silver staining of conditioned media provided in Fig.1E. **B,** Table of identified proteins under static or stretch condition.

| | | | | Ratio | | | T-Test | | |
|---|---|---|---|---|---|---|---|---|---|
| No | Symbol | Name | Num. of Peptide | Stretch/Static | Stretch/Stretch+BFA | Stretch+BFA/Static | Stretch/Static | Stretch/Stretch+BFA | Stretch+BFA/Static |
| 1 | Lum | Lumican | 13 | 7.64 | 21.42 | 0.36 | 2.09E-05 | 2.84E-04 | 5.06E-05 |
| 2 | Thbs1 | Thrombospondin 1 | 42 | 5.80 | 13.40 | 0.43 | 6.23E-06 | 1.00E-04 | 1.84E-04 |
| 3 | Col11a1 | Collagen alpha-1(XI) chain | 15 | 4.88 | 6.53 | 0.75 | 1.59E-05 | 1.17E-04 | 6.49E-02 |
| 4 | Col3A1 | Collagen alpha-1(III) chain | 211 | 4.85 | 15.08 | 0.32 | 8.83E-06 | 6.83E-05 | 9.59E-04 |
| 5 | Postn | Periostin | 17 | 4.75 | 6.89 | 0.69 | 2.60E-04 | 1.93E-03 | 2.53E-03 |
| 6 | EPYC | Epiphycan | 18 | 4.70 | 20.24 | 0.23 | 1.70E-05 | 1.86E-04 | 1.46E-06 |
| 7 | Fbn1 | Fibrillin 1 | 108 | 4.67 | 10.21 | 0.46 | 5.65E-06 | 7.29E-05 | 2.97E-04 |
| 8 | Mmp2 | Matrix Metalloproteinase-2 | 42 | 4.65 | 16.28 | 0.29 | 1.18E-04 | 8.20E-04 | 2.16E-06 |
| 9 | Bgn | Biglycan | 44 | 4.61 | 36.24 | 0.13 | 1.57E-04 | 6.72E-04 | 1.50E-03 |
| 10 | Col6a2 | Collagen type VI alpha 2 chain | 12 | 4.54 | 6.20 | 0.73 | 4.48E-03 | 1.69E-02 | 1.56E-02 |
| 11 | Col4a2 | Collagen type IV alpha 2 chain | 22 | 4.46 | 9.77 | 0.46 | 1.39E-04 | 1.02E-03 | 3.39E-06 |
| 12 | Ogn | Osteoglycin | 8 | 4.43 | 8.18 | 0.54 | 2.69E-04 | 1.60E-03 | 3.89E-03 |
| 13 | Nid2 | Nidogen-2 | 34 | 4.39 | 6.39 | 0.69 | 1.96E-04 | 1.22E-03 | 4.56E-02 |
| 14 | Col5a1 | Collagen alpha-1(V) chain | 29 | 4.28 | 7.35 | 0.58 | 2.18E-05 | 2.23E-04 | 2.53E-03 |
| 15 | Serpinf2 | Alpha-2 antiplasmin | 18 | 4.27 | 14.51 | 0.29 | 9.79E-03 | 2.17E-02 | 1.87E-03 |
| 16 | Ltbp2 | Latent-transforming growth factor beta-binding protein 2 | 21 | 4.19 | 6.11 | 0.69 | 5.12E-06 | 9.69E-05 | 3.42E-05 |
| 17 | Timp1 | Metalloproteinase inhibitor 1 | 16 | 4.19 | 12.93 | 0.32 | 1.87E-05 | 1.16E-04 | 7.80E-04 |
| 18 | Fn1 | Fibronectin | 300 | 4.17 | 7.19 | 0.58 | 1.10E-04 | 8.98E-04 | 2.04E-03 |
| 19 | Vcan | Versican core protein (Fragments) | 10 | 4.11 | 4.80 | 0.86 | 3.14E-04 | 2.05E-03 | 2.23E-01 |
| 20 | Igfbp7 | Insulin-like growth factor binding protein 7, isoform CRA_b | 26 | 4.05 | 28.38 | 0.14 | 1.20E-04 | 5.67E-04 | 2.48E-04 |
| 21 | B2m | Beta-2-microglobulin | 5 | 3.97 | 6.62 | 0.60 | 5.97E-05 | 5.55E-04 | 7.83E-04 |
| 22 | Lox | Lysyl Oxidase | 4 | 3.93 | 5.25 | 0.75 | 5.52E-04 | 4.71E-04 | 3.74E-01 |
| 23 | Sparc | Secreted protein acidic and cysteine rich | 58 | 3.92 | 32.51 | 0.12 | 1.03E-04 | 5.40E-04 | 2.64E-05 |
| 24 | Cxcl1 | Growth-regulated alpha protein | 5 | 3.90 | 16.64 | 0.23 | 8.53E-05 | 2.85E-04 | 2.27E-03 |
| 25 | C1r | Complement C1r | 37 | 3.84 | 17.04 | 0.23 | 5.62E-05 | 3.70E-04 | 1.47E-05 |
| 26 | Col1a2 | Collagen alpha-2(I) chain | 438 | 3.84 | 17.30 | 0.22 | 5.28E-08 | 3.98E-07 | 4.45E-06 |
| 27 | Hspg2 | Heparan Sulfate proteoglycan 2 | 67 | 3.80 | 6.15 | 0.62 | 3.53E-03 | 1.17E-02 | 1.31E-02 |
| 28 | Col4a1 | Collagen type IV alpha 1 chain | 8 | 3.79 | 17.94 | 0.21 | 1.37E-05 | 1.24E-04 | 4.27E-06 |
| 29 | Sod1 | Superoxide dismutase 1 | 18 | 3.73 | 22.80 | 0.16 | 2.42E-04 | 9.24E-04 | 3.54E-04 |
| 30 | Timp2 | Metalloproteinase inhibitor 2 | 17 | 3.71 | 16.34 | 0.23 | 3.83E-05 | 2.76E-04 | 2.82E-06 |
| 31 | Fstl1 | Follistatin-related protein 1 | 34 | 3.70 | 30.34 | 0.12 | 2.65E-04 | 1.10E-03 | 3.14E-06 |
| 32 | Col1a1 | Collagen alpha-1(I) chain | 586 | 3.67 | 10.71 | 0.34 | 8.08E-06 | 6.12E-05 | 2.44E-04 |
| 33 | Cdh13 | Cadherin 13 | 3 | 3.66 | 3.81 | 0.96 | 6.36E-05 | 8.08E-04 | 1.76E-01 |
| 34 | C1s | Complement C1s subcomponent | 9 | 3.57 | 15.26 | 0.23 | 2.97E-05 | 2.16E-04 | 1.89E-05 |
| 35 | Igfbp6 | Insulin-like growth factor-binding protein 6 | 7 | 3.56 | 13.33 | 0.27 | 2.33E-04 | 3.81E-04 | 7.42E-03 |
| 36 | Fbln2 | Fibulin 2 | 43 | 3.54 | 4.16 | 0.85 | 3.61E-06 | 1.22E-04 | 1.54E-01 |
| 37 | Ecm1 | Extracellular matrix protein 1 | 12 | 3.54 | 12.17 | 0.29 | 7.76E-06 | 3.12E-05 | 3.04E-04 |
| 38 | Col6a1 | Collagen type VI alpha 1 chain | 12 | 3.53 | 5.49 | 0.64 | 2.16E-04 | 1.44E-03 | 2.31E-04 |
| 39 | Tcn2 | Transcobalamin-2 | 12 | 3.51 | 10.61 | 0.33 | 3.29E-04 | 1.47E-03 | 1.52E-05 |
| 40 | Il1rl1 | Interleukin-1 receptor-like 1 | 3 | 3.50 | 4.10 | 0.85 | 4.58E-04 | 2.77E-03 | 1.33E-01 |
| 41 | Rnase4 | Ribonuclease 4 | 3 | 3.45 | 10.26 | 0.34 | 4.78E-05 | 3.14E-04 | 4.12E-05 |
| 42 | Spp1 | Osteopontin | 22 | 3.40 | 15.29 | 0.22 | 5.10E-03 | 1.07E-02 | 7.08E-06 |
| 43 | Inhba | Inhibin beta A chain | 6 | 3.36 | 5.91 | 0.57 | 1.29E-04 | 6.44E-04 | 4.20E-03 |
| 44 | Loxl3 | Lysyl oxidase-like 3 | 7 | 3.35 | 5.04 | 0.67 | 1.36E-06 | 1.99E-05 | 2.62E-04 |
| 45 | Eln | Elastin | 13 | 3.35 | 3.35 | 1.00 | 6.17E-04 | 4.80E-03 | #DIV/0! |
| 46 | Lamb1 | Laminin subunit beta 1 | 14 | 3.28 | 3.57 | 0.92 | 2.23E-03 | 1.10E-02 | 1.63E-01 |
| 47 | Nucb1 | Nucleobindin-1 | 27 | 3.27 | 10.99 | 0.30 | 7.79E-06 | 7.26E-05 | 7.89E-06 |
| 48 | Plod2 | Procollagen-lysine,2-oxoglutarate 5-dioxygenase 2 | 5 | 3.25 | 3.79 | 0.86 | 6.74E-03 | 2.29E-02 | 5.06E-01 |
| 49 | Pcolce | Procollagen C-endopeptidase enhancer 1 | 37 | 3.24 | 17.52 | 0.19 | 5.76E-05 | 3.02E-04 | 7.95E-06 |
| 50 | Clstn1 | Calsyntenin-1 | 24 | 3.21 | 9.28 | 0.35 | 1.68E-05 | 9.09E-05 | 1.80E-04 |
| 51 | Col5a2 | Collagen type V alpha 2 chain | 69 | 3.19 | 8.69 | 0.37 | 2.94E-05 | 2.31E-04 | 2.05E-06 |
| 52 | Ccl2 | C-C motif chemokine 2 | 5 | 3.18 | 14.82 | 0.21 | 4.42E-04 | 1.10E-03 | 1.01E-03 |
| 53 | Svep1 | Sushi, von Willebrand factor type A, EGF and pentraxin domain-containing protein 1 | 14 | 3.17 | 3.96 | 0.80 | 3.34E-05 | 3.75E-04 | 2.22E-03 |
| 54 | Srpx2 | Sushi repeat-containing protein SRPX2 | 4 | 3.15 | 4.81 | 0.65 | 1.09E-05 | 1.35E-04 | 8.35E-05 |
| 55 | Pin1 | Peptidylprolyl isomerase | 8 | 3.11 | 5.42 | 0.57 | 1.42E-04 | 5.34E-04 | 1.32E-02 |
| 56 | Clu | Clusterin | 24 | 3.11 | 18.06 | 0.17 | 9.96E-05 | 3.74E-04 | 6.98E-05 |
| 57 | Lamc1 | Laminin subunit gamma 1 | 24 | 3.09 | 3.83 | 0.81 | 2.01E-03 | 8.77E-03 | 1.11E-02 |
| 58 | Pxdn | Peroxidasin | 3 | 3.09 | 3.99 | 0.77 | 3.19E-03 | 1.16E-02 | 2.80E-02 |
| 59 | Nid1 | Nidogen-1 (Fragment) | 8 | 3.06 | 8.55 | 0.36 | 1.59E-04 | 7.57E-04 | 9.59E-06 |
| 60 | Atp5b | ATP synthase subunit beta, mitochondrial | 3 | 3.04 | 3.44 | 0.88 | 8.36E-04 | 3.97E-03 | 3.91E-01 |
| 61 | Lgals3bp | Galectin-3-binding protein | 21 | 2.99 | 11.79 | 0.25 | 5.39E-04 | 7.44E-04 | 4.46E-03 |
| 62 | Htra1 | Serine protease HTRA1 | 20 | 2.94 | 5.32 | 0.55 | 2.96E-04 | 1.47E-03 | 4.64E-04 |
| 63 | Ltbp1 | Latent-transforming growth factor beta-binding protein 1 | 8 | 2.93 | 3.32 | 0.88 | 2.96E-04 | 1.91E-03 | 1.18E-01 |
| 64 | Acan | Aggrecan core protein | 7 | 2.93 | 2.94 | 1.00 | 1.24E-03 | 8.02E-03 | 3.74E-01 |
| 65 | Olfml3 | Olfactomedin-like protein 3 | 6 | 2.92 | 2.73 | 1.07 | 1.45E-03 | 1.84E-03 | 8.41E-01 |
| 66 | Cpe | Carboxypeptidase E | 7 | 2.91 | 5.33 | 0.55 | 8.69E-04 | 3.19E-03 | 3.12E-04 |
| 67 | Oaf | Out at first protein homolog | 3 | 2.91 | 3.21 | 0.91 | 1.90E-04 | 1.58E-03 | 6.06E-02 |
| 68 | Col8a1 | Collagen type VIII alpha 1 chain | 11 | 2.90 | 2.82 | 1.03 | 1.10E-03 | 8.17E-03 | 6.39E-01 |
| 69 | Nov | Protein NOV homolog | 4 | 2.77 | 6.47 | 0.43 | 4.14E-04 | 1.38E-03 | 2.66E-04 |
| 70 | Col18a1 | Collagen type XVIII alpha 1 chain | 17 | 2.75 | 7.72 | 0.36 | 3.29E-04 | 1.11E-03 | 6.31E-05 |
| 71 | Cst3 | Cystatin-C | 9 | 2.70 | 10.90 | 0.25 | 9.19E-04 | 8.24E-05 | 1.48E-02 |
| 72 | Fbln5 | Fibulin-5 | 9 | 2.69 | 6.68 | 0.40 | 4.91E-04 | 1.67E-03 | 9.44E-07 |
| 73 | Syncrip | Heterogeneous nuclear ribonucleoprotein Q | 3 | 2.67 | 2.42 | 1.10 | 6.05E-03 | 6.63E-06 | 8.51E-01 |
| 74 | Ctsb | Cathepsin B | 41 | 2.63 | 5.35 | 0.49 | 6.15E-05 | 6.22E-05 | 2.88E-03 |
| 75 | Minpp1 | Multiple inositol polyphosphate phosphatase 1 | 3 | 2.53 | 2.68 | 0.95 | 4.47E-03 | 1.89E-02 | 1.54E-02 |
| 76 | Csf1 | Macrophage colony-stimulating factor 1 | 9 | 2.53 | 7.14 | 0.35 | 4.08E-04 | 1.13E-03 | 6.14E-05 |
| 77 | Ctsl | Cathepsin L1 | 10 | 2.50 | 5.81 | 0.43 | 3.54E-05 | 3.64E-05 | 8.06E-04 |
| 78 | Efemp2 | EGF-containing fibulin extracellular matrix protein 2 | 8 | 2.47 | 4.63 | 0.53 | 1.67E-03 | 3.76E-03 | 2.84E-03 |
| 79 | C3 | Complement C3 | 20 | 2.27 | 3.47 | 0.65 | 3.84E-05 | 4.87E-05 | 3.00E-03 |
| 80 | Pai1 | Plasminogen activator inhibitor 1 | 16 | 2.27 | 6.86 | 0.33 | 5.37E-03 | 6.78E-03 | 3.70E-04 |
| 81 | Npc2 | NPC intracellular cholesterol transporter 2 | 8 | 2.24 | 2.41 | 0.93 | 2.15E-04 | 1.53E-03 | 1.30E-01 |
| 82 | Ctgf | Connective tissue growth factor | 13 | 2.19 | 6.79 | 0.32 | 7.35E-04 | 1.07E-03 | 3.50E-04 |
| 83 | Pdgfrl | Platelet-derived growth factor receptor-like protein | 3 | 2.12 | 2.21 | 0.96 | 1.45E-05 | 1.65E-05 | 3.74E-01 |
| 84 | Rps8 | 40S ribosomal protein S8 | 6 | 0.48 | 0.42 | 1.14 | 6.40E-04 | 3.59E-05 | 1.39E-01 |
| 85 | Rps16 | 40S ribosomal protein S16 | 5 | 0.35 | 0.29 | 1.21 | 4.91E-03 | 3.72E-02 | 4.24E-01 |
| 86 | Rps26 | 40S ribosomal protein S26 | 3 | 0.41 | 0.78 | 0.53 | 9.30E-04 | 8.81E-01 | 2.56E-02 |
| 87 | Rpl30 | 60S ribosomal protein L30 | 4 | 0.41 | 0.38 | 1.07 | 1.56E-03 | 8.52E-03 | 5.85E-01 |

**Online Figure II. List of identified secreted proteins from rat vascular SMCs under cyclic stretch.**

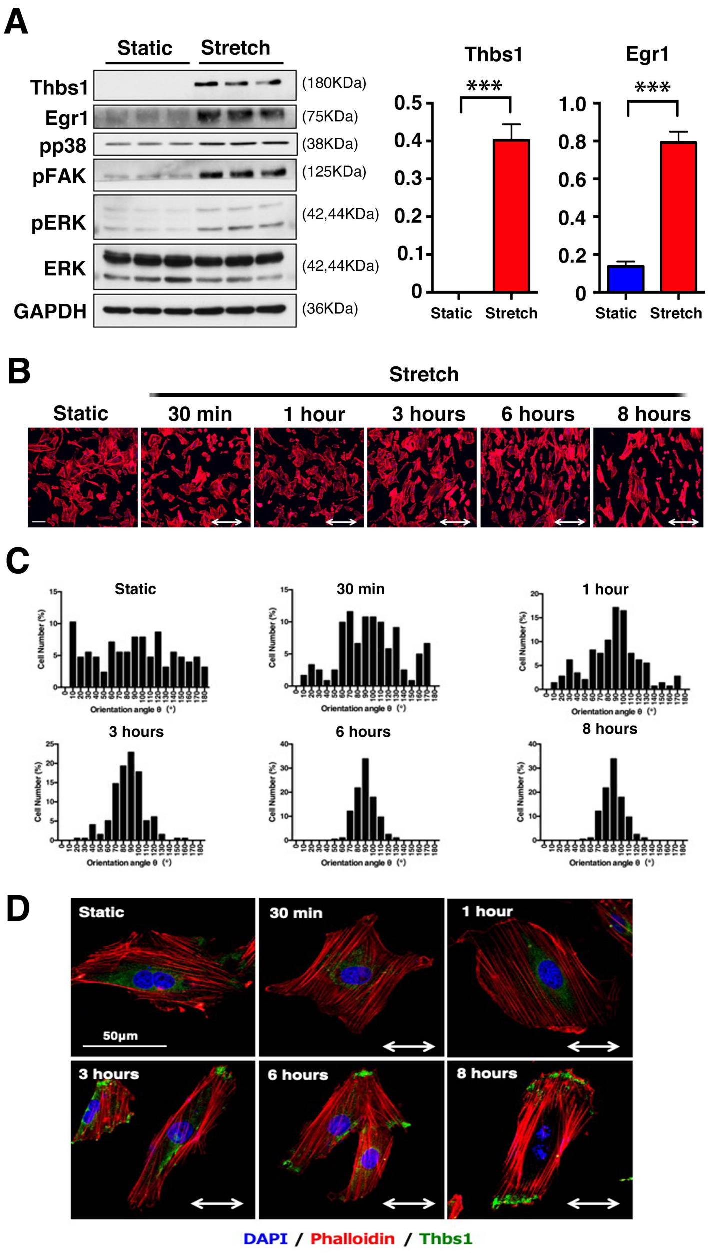


**Online Figure III. Thbs1 expression during cyclic stretch.** Rat vascular SMCs were subjected to cyclic stretch (1.0 Hz, 20% strain, 8h). **A,** Cell lysates were analyzed using Western blotting with indicated antibodies (n=3). Quantification graphs are shown on the right. Bars are means ± SEM. ***$P$<0.001, unpaired $t$-test. **B-C,** Immunostaining with Phalloidin (red). Cell orientation was analyzed by measuring the orientation angle (θ) of the long axis of the ellipse relative to the stretch axis. 50-100 cells were evaluated in each condition (n=3). Bars are 50 μm. Two-way arrows indicates stretch direction. **D,** Representative immunostaining of SMCs at an indicated time after stretch. Phalloidin (red), Thbs1 (green), and DAPI (blue). n=3.

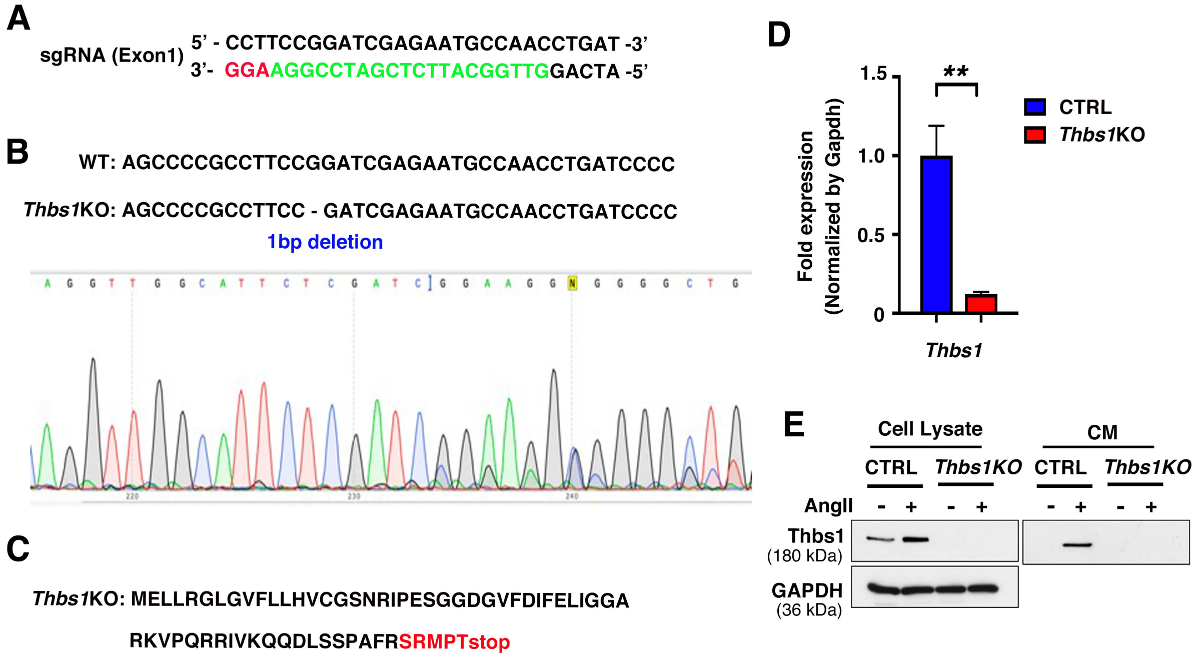


**Online Figure IV.** ***Thbs1* deletion by CRISPR-Cas9 in rat vascular SMCs. A,** sgRNA sequence (green) targeting exon1 of *Thbs1*. PAM (protospacer adjacent motif) sequence is shown in red. **B,** Sequence comparison between *Thbs1*KO cell line (*Thbs1*KO) and control (CTRL) cell line. **C,** Amino acid sequence for *Thbs1*KO. The altered amino acid sequences are shown in red and stop codon appeared in Exon 1 of Thbs1 due to a frame shift (1bp deletion). **D,** qPCR analysis confirming the efficiency of gene deletion in rat vascular SMCs. Bars are means ± SEM. ***P* < 0.01, unpaired *t*-test. **E,** CTRL and *Thbs1*KO cells cultured in serum-free media for 24 h, then stimulated with Ang II (100 nM) for 6 h. Cell lysates and conditioned medium (CM) were analyzed using Western blotting with indicated antibodies. n=3.

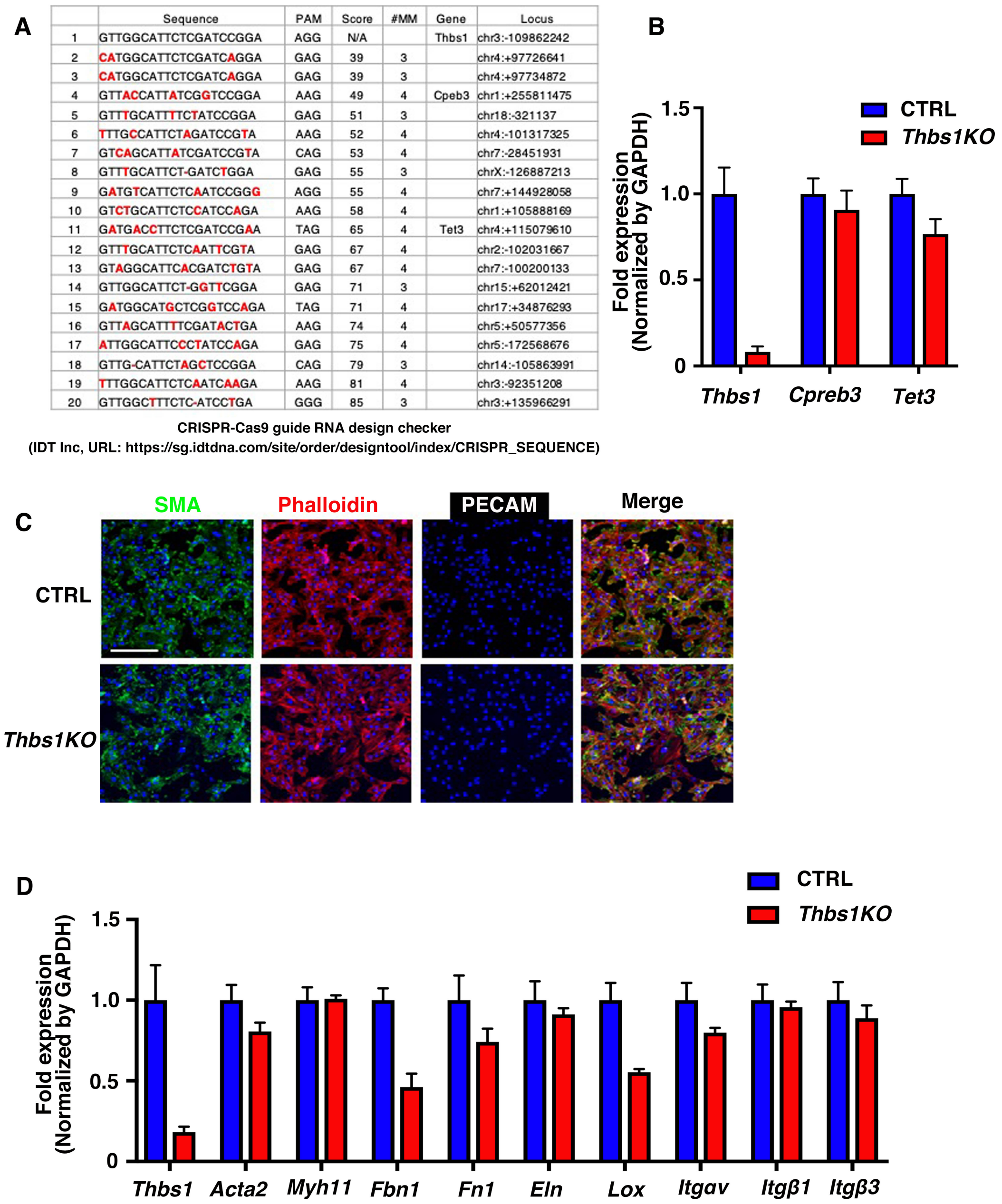

A

| | Sequence | PAM | Score | #MM | Gene | Locus |
|---|---|---|---|---|---|---|
| 1 | GTTGGCATTCTCGATCCGGA | AGG | N/A | | Thbs1 | chr3:-109862242 |
| 2 | CATGGCATTCTCGATCAGGA | GAG | 39 | 3 | | chr4:+97726641 |
| 3 | CATGGCATTCTCGATCAGGA | GAG | 39 | 3 | | chr4:+97734872 |
| 4 | GTTACCATTATCGGTCCGGA | AAG | 49 | 4 | Cpeb3 | chr1:+255811475 |
| 5 | GTTTGCATTTTCTATCCGGA | GAG | 51 | 3 | | chr18:-321137 |
| 6 | TTTGCCATTCTAGATCCGTA | AAG | 52 | 4 | | chr4:-101317325 |
| 7 | GTCAGCATTATCGATCCGTA | CAG | 53 | 4 | | chr7:-28451931 |
| 8 | GTTTGCATTCT-GATCTGGA | GAG | 55 | 3 | | chrX:-126887213 |
| 9 | GATGTCATTCTCAATCCGGG | AGG | 55 | 4 | | chr7:+144928058 |
| 10 | GTCTGCATTCTCCATCCAGA | AAG | 58 | 4 | | chr1:+105888169 |
| 11 | GATGACCTTCTCGATCCGAA | TAG | 65 | 4 | Tet3 | chr4:+115079610 |
| 12 | GTTTGCATTCTCAATTCGTA | GAG | 67 | 4 | | chr2:-102031667 |
| 13 | GTAGGCATTCACGATCTGTA | GAG | 67 | 4 | | chr7:-100200133 |
| 14 | GTTGGCATTCT-GGTTCGGA | GAG | 71 | 3 | | chr15:+62012421 |
| 15 | GATGGCATGCTCGGTCCAGA | TAG | 71 | 4 | | chr17:+34876293 |
| 16 | GTTAGCATTTTCGATACTGA | AAG | 74 | 4 | | chr5:+50577356 |
| 17 | ATTGGCATTCCCTATCCAGA | GAG | 75 | 4 | | chr5:-172568676 |
| 18 | GTTG-CATTCTAGCTCCGGA | CAG | 79 | 3 | | chr14:-105863991 |
| 19 | TTTGGCATTCTCAATCAAGA | AAG | 81 | 4 | | chr3:-92351208 |
| 20 | GTTGGCTTTCTC-ATCCTGA | GGG | 85 | 3 | | chr3:+135966291 |

CRISPR-Cas9 guide RNA design checker
(IDT Inc, URL: https://sg.idtdna.com/site/order/designtool/index/CRISPR_SEQUENCE)



**Online Figure V. Off-target effects on *Thbs1 KO* cells. A,** Potential off target list of sgRNA for Thbs1. Red indicates miss march (#MM) nucleotides. **B,** qPCR analysis confirming the expression of *Creb3* and *Tet3* in CTRL and *Thbs1KO* rat vascular SMCs. Bars indicate technical triplicate. **C,** Immunostaining of CTRL or *Thbs1KO* cells with smooth muscle actin (SMA, green), Phalloidin (red) and PECAM (white). Bar is 50 µm. **D,** qPCR analysis for the expression of SMC contractile marker (*Acta2* and *Myh11)*, matrix compound (*Fbn1*, *Fn1*, *Eln* and *Lox*) and integrins (*Itgαv*, *Itgβ1* and *Itgβ3*) in CTRL and *Thbs1KO* rat vascular SMCs. Bars indicate technical triplicate.

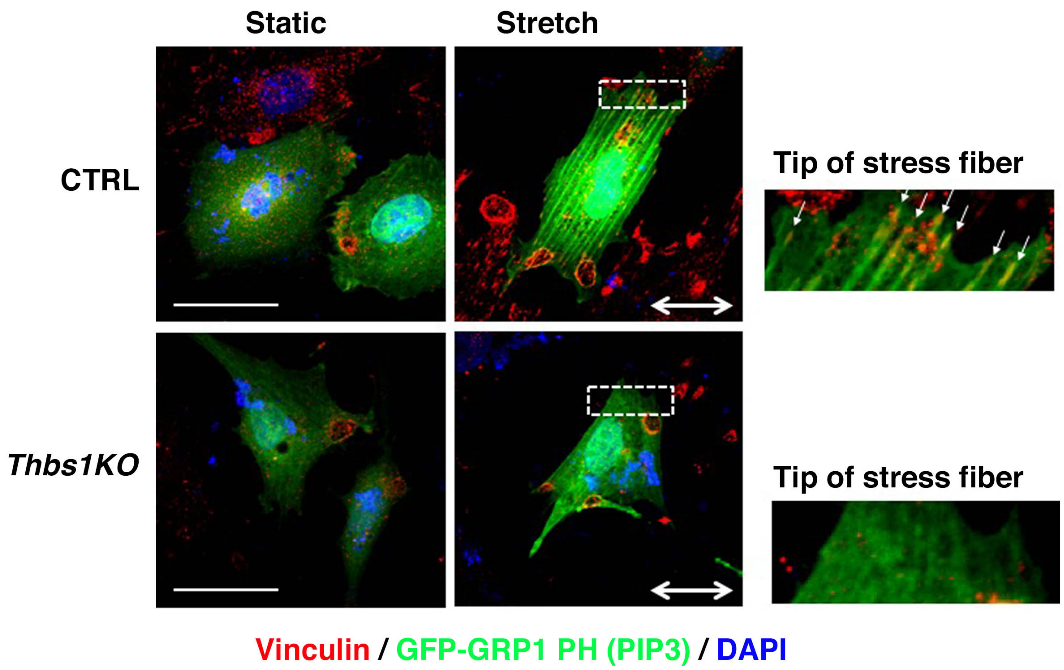


**Online Figure VI. Deletion of *Thbs1* affects maturation of FA.** Representative Immunostaining of control (CTRL) or *Thbs1*KO cells with vinculin (red) in static or stretch condition. GFP-GRP1-PH (green) and DAPI (blue). White arrows show vinculin deposition onto the tip of actin fibers. (n=3). Two-way arrows indicate stretch direction. Bars are 50 μm.

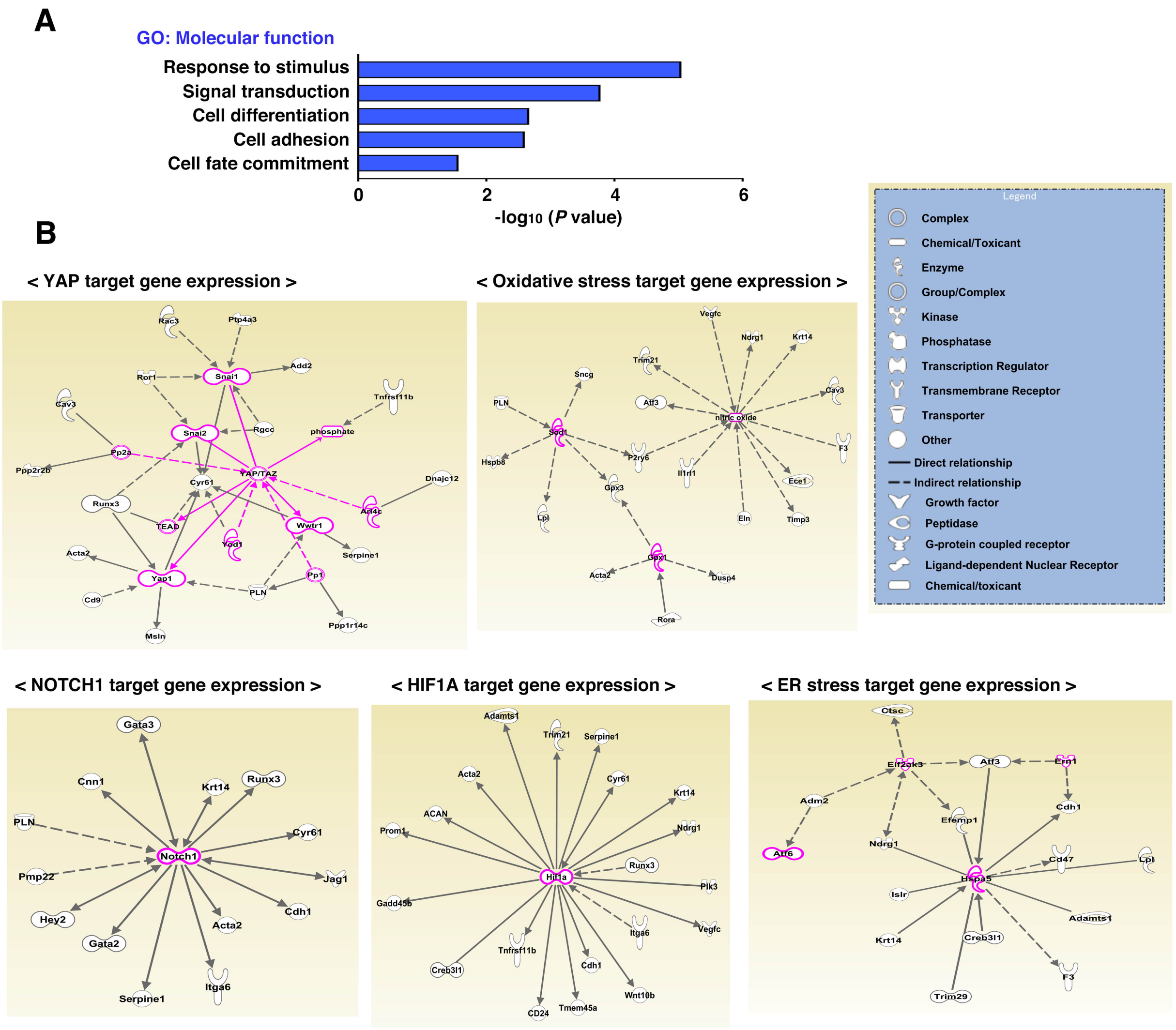


**Online Figure VII. Bioinformatic analysis for RNA-seq. A,** Functional enrichment analysis of 289 genes in Fig. 4C-D. Top enriched GO terms associated with molecular functions (blue) are shown. **B,** Among 289 genes in Fig. 4C-D, network analysis was performed using YAP, oxidative stress-related gene-, NOTCH1, HIF1A, and ER stress-related gene categories via IPA. Diagrams show direct (solid lines) and indirect (dashed lines) interactions among genes.

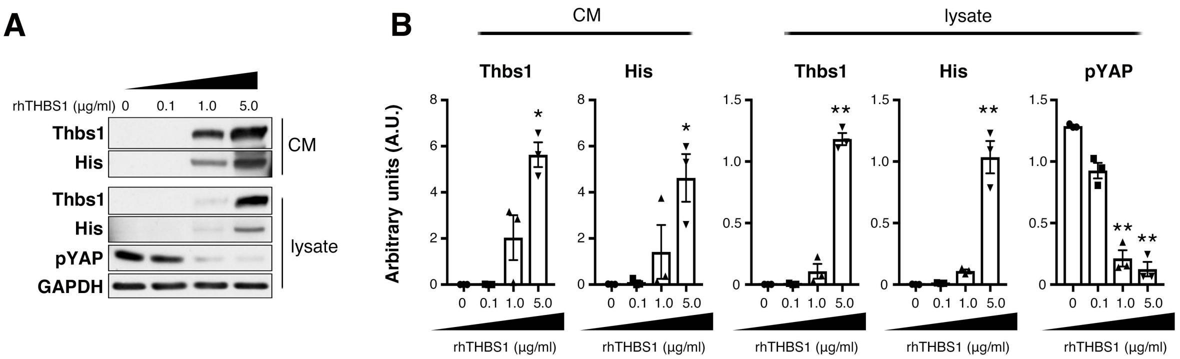


**Online Figure VIII. Add-back experiment in *Thbs1KO* cells upon cyclic stretch. A,** *Thbs1KO* rat SMCs were treated with or without recombinant human THBS1 with His-tag (rhThbs1;0.1 or 1.0 or 5.0 μg/ml) followed by 24 h of serum starvation and then subjected to uniaxial cyclic stretch (20% strain; 1Hz) for 8h. Representative Western blots shows Thbs1 and His in conditioned medium (CM), Thbs1, His and pYAP levels in cell lysate (n=3). **B,** Quantification graphs shown in right. Bars are means ± SEM. **P*<0.05, ***P*<0.01, one-way ANOVA.

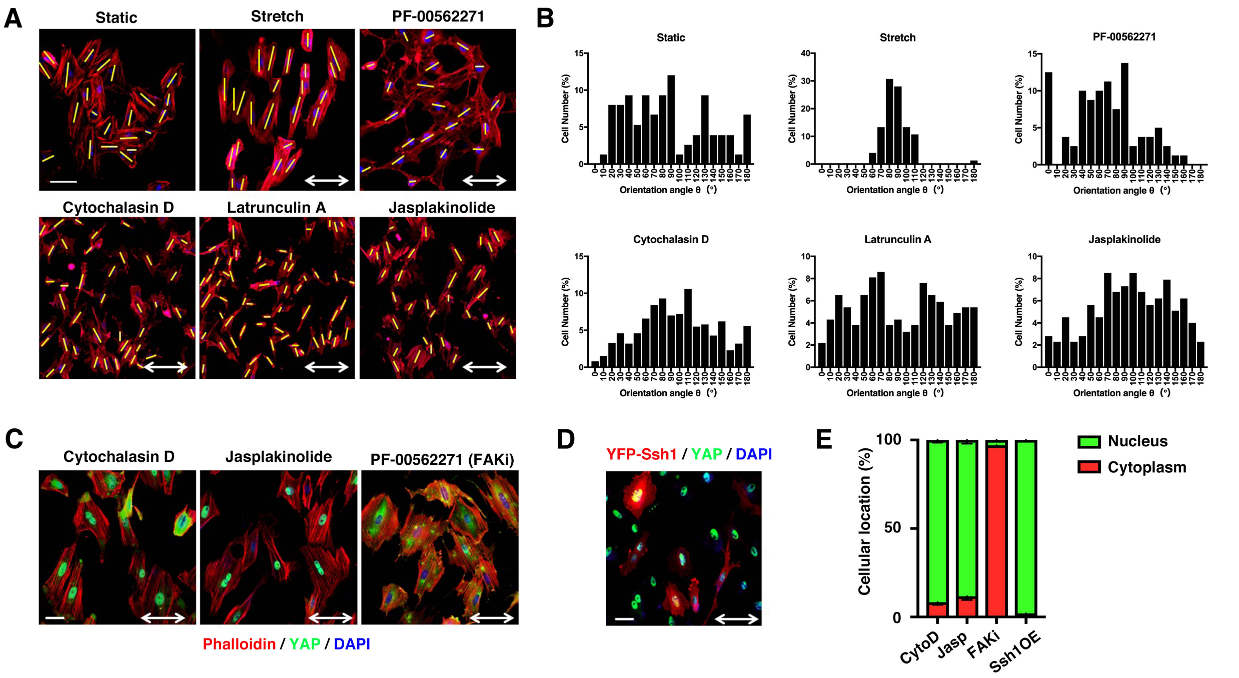


**Online Figure IX. FA and actin remodeling are required for proper cell orientation in response to cyclic stretch.** Rat vascular SMCs were cultured in serum-free media for 24 h and treated with Cytochalasin D (1 μM), Latrunclin A (1 μM), Jasplakinolide (20 nM) or PF-00562271 (100 nM, FAK inhibitor) and subjected to cyclic stretch (1.0 Hz, 20 % strain, 8h). Bars are 50 μm. Two-way arrows indicate stretch direction. **A,** Immunostaining with Phalloidin (red). **B,** Cell orientation was analyzed by measuring the orientation angle (θ) of the long axis (yellow bars in a) of the ellipse relative to the stretch axis. 50-100 cells were counted in each condition (n=3). **C,** Rat vascular SMCs were cultured in serum-free media for 24 h and treated with Cytochalasin D (1 μM), Jasplakinolide (20 nM), or FAK inhibitor (100 nM) during cyclic stretch (n=3). Representative immunostaining with YAP (green), Phalloidin (red), and DAPI (blue). Two-way arrows indicate stretch direction. Bars are 50 μm. **D,** Rat vascular SMCs were transfected with YFP-Ssh1 (red) and cyclic stretch was performed at 48 h after transfection (n=3). Representative immunostaining with YAP (green) and DAPI (blue). Two-way arrows indicate stretch direction. Bars are 50 μm. **E,** Quantification of YAP localization in C and D. 50-150 cells were evaluated in each experiment (n=3).

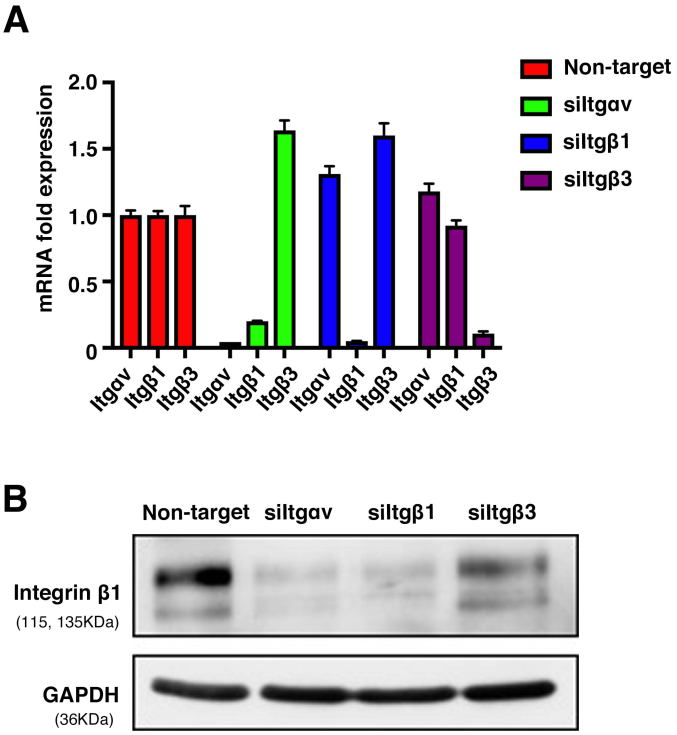


**Online Figure X. Deletion of integrins by siRNA. A,** Quantitative polymerase chain reaction (qPCR) for confirmation of knockdown of *Itgαv*, *Itgβ1* and *Itgβ3*. Total RNA from small interfering RNA (siRNA)-treated cells was extracted 2 days after transfection. The fold expression relative to scramble siRNA (non-target) is shown. Bars indicate technical triplicate. **B,** Western blot shows integrin β1 levels in siRNA-treated cells.

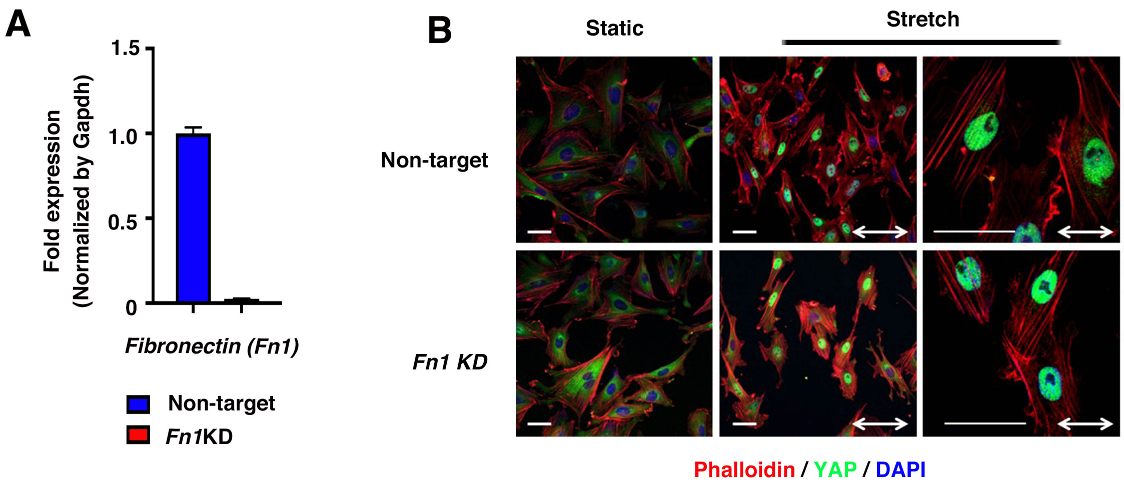


**Online Figure XI. Knockdown of fibronectin does not affect stretch-induced nuclear shuttling of YAP. A,** qPCR analysis confirming the efficiency of *Fn1* knockdown in rat vascular SMCs. Bars indicate technical triplicate. **B,** Non-target or *Fn1* siRNA-treated rat vascular SMCs were subjected to cyclic stretch (n=3). Immunostaining with YAP (green). Phalloidin (red) and DAPI (blue). Bars are 50 μm.

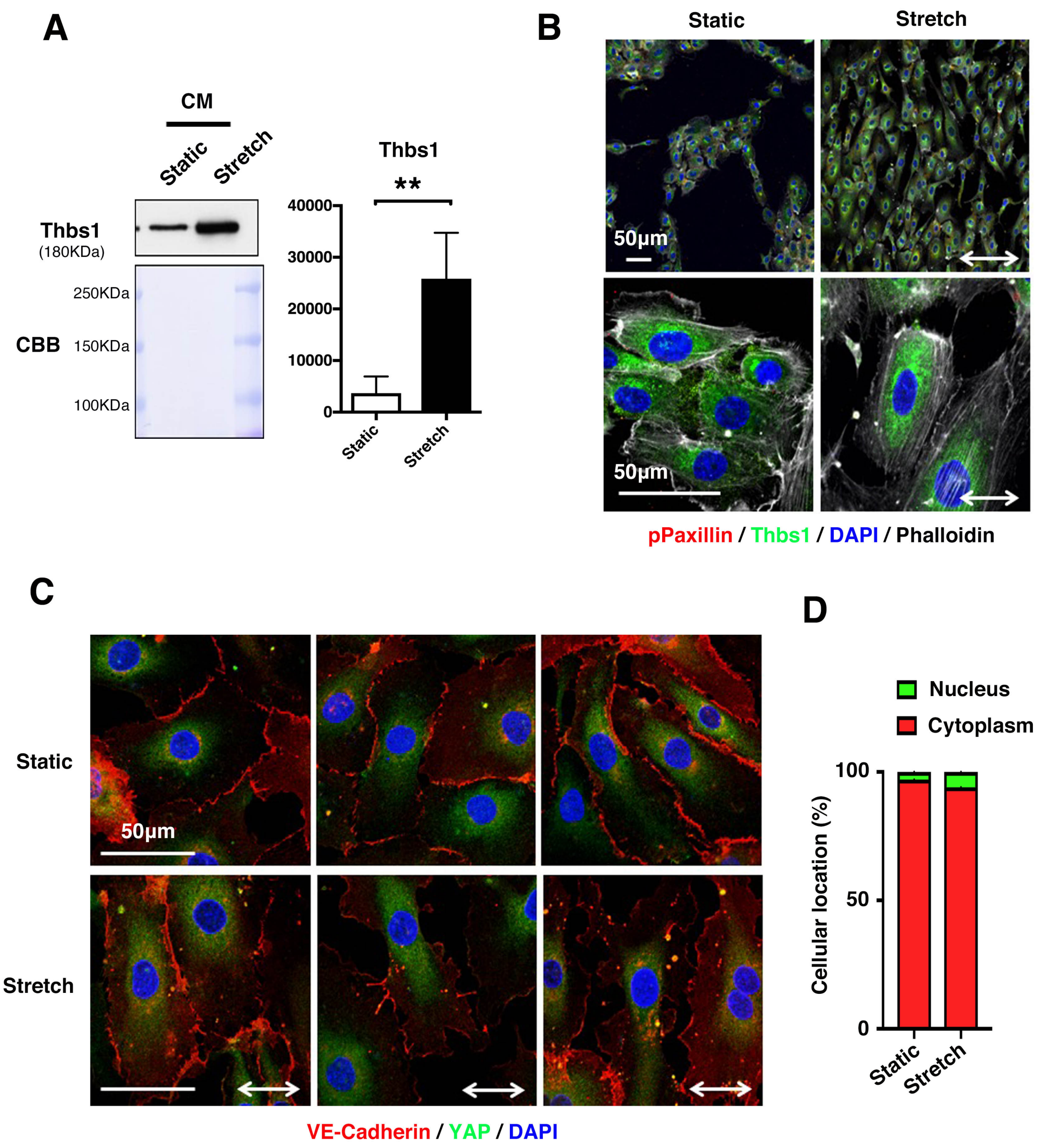


**Online Figure XII. Cyclic stretch did not induce translocation of Thbs1 to FAs and nuclear shuttling of YAP in HUVECs. A,** Representative Western blot for Thbs1 using the conditioned medium (CM) of HUVEC (n=3) in static or stretch condition. Quantification graph is shown in right. $^{**}P < 0.01$, unpaired *t*-test. **B,** HUVECs were subjected to cyclic stretch (1.0 Hz, 20% strain, 8 h). Bars are 50 μm. Two-way arrows indicate stretch direction. Immunostaining with p-Paxillin (red), Thbs1 (green), Phalloidin (grey) and DAPI (blue). **C,** Representative immunostaining of HUVEC in static or stretch condition with VE-cadherin (red), YAP (green), and DAPI (blue). **D,** Quantification of YAP localization in 80-100 cells were evaluated in each condition (n=3). YAP localizes in nuclear (green) or cytoplasm (red).

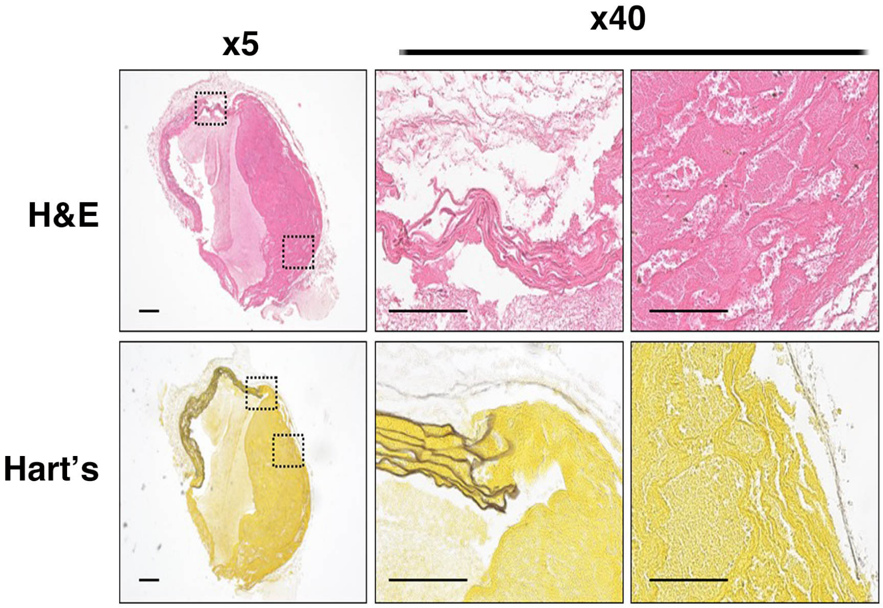


**Online Figure XIII. Representative image of *Thbs1* null (*Thbs1KO*) mice with aortic rupture after TAC.** H&E and Hart's staining of the ascending aortas in TAC-operated *Thbs1KO* mice. 7 mice died after TAC in *Thbs1KO* mice (Fig. 6B) and 3 of 7 mice showed aortic rupture. Bars are 100 μm.

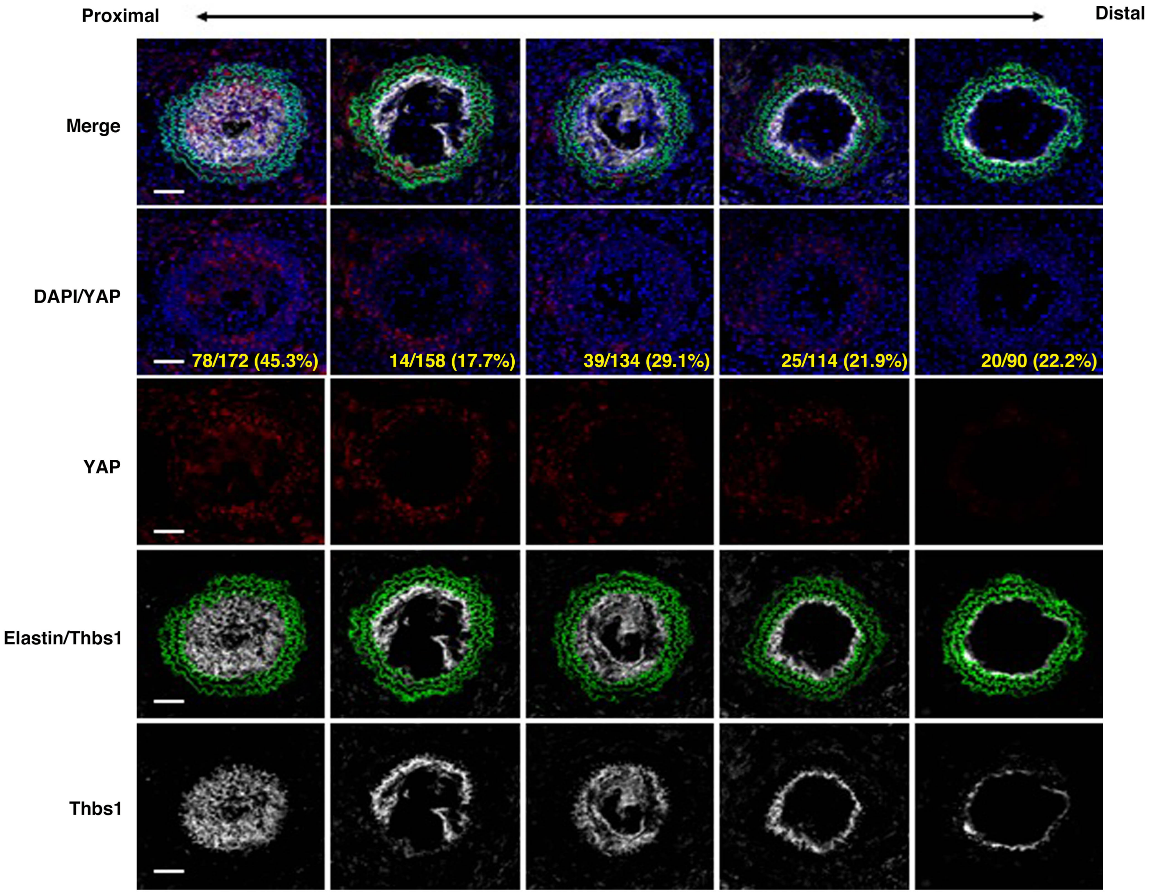


**Online Figure XIV. Neointima formation at 1 week after carotid artery ligation.** Representative immunostaining of cross sections of ligated arteries with YAP (red) and Thbs1 (white) at 1 week after ligation in WT mice (n=5). DAPI (blue) and auto-fluorescence of elastin (green). Images shows are shown in proximal to distal direction from the ligature point. Bars are 50 μm. Quantification of co-localization with YAP and DAPI using Imaris colocalization software were indicated by yellow (number of nuclear YAP / total number of nucleus).

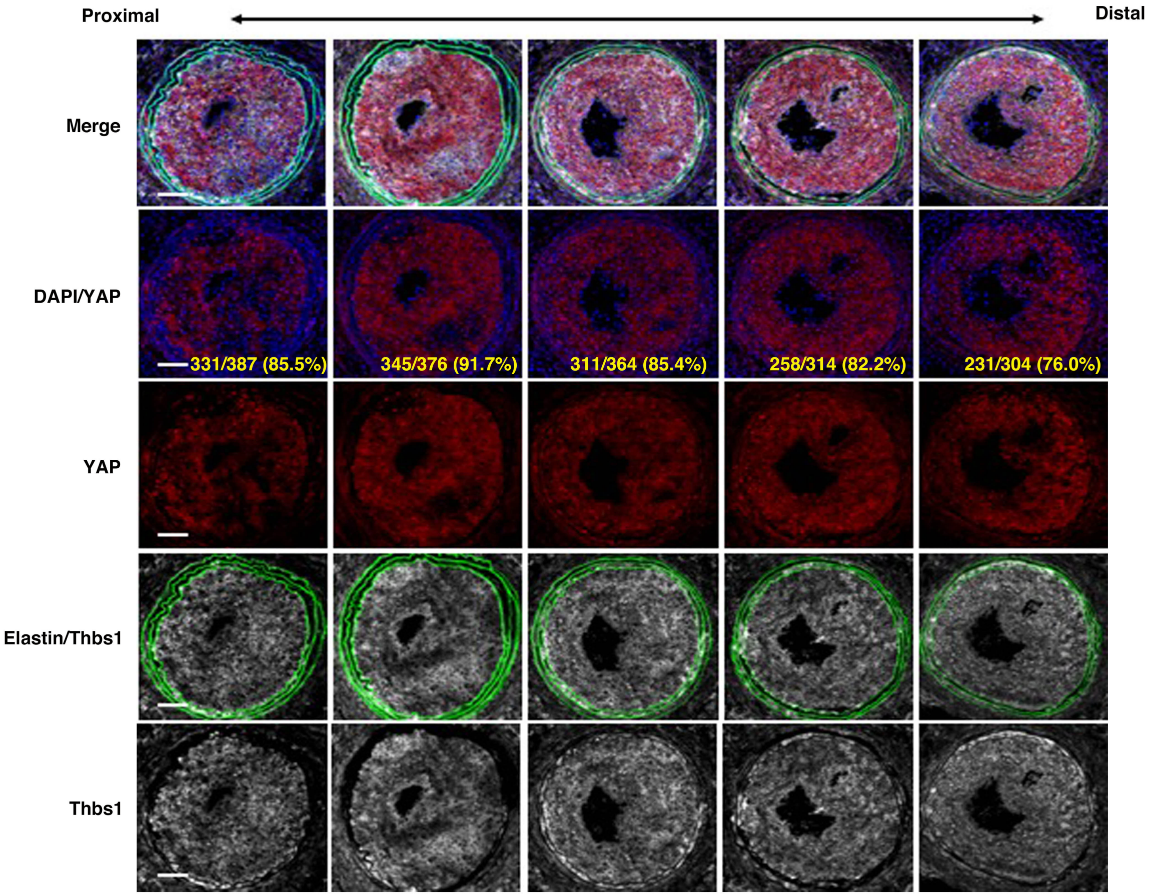


**Online Figure XV. Neointima formation at 2 weeks after carotid artery ligation.** Representative immunostaining of cross sections of ligated arteries with YAP (red) and Thbs1 (white) at 2 weeks after ligation in WT mice (n=5). DAPI (blue) and auto-fluorescence of elastin (green). Images shows are shown in proximal to distal direction from the ligature point. Bars are 50 μm. Quantification of co-localization with YAP and DAPI using Imaris colocalization software were indicated by yellow (number of nuclear YAP / total number of nucleus).

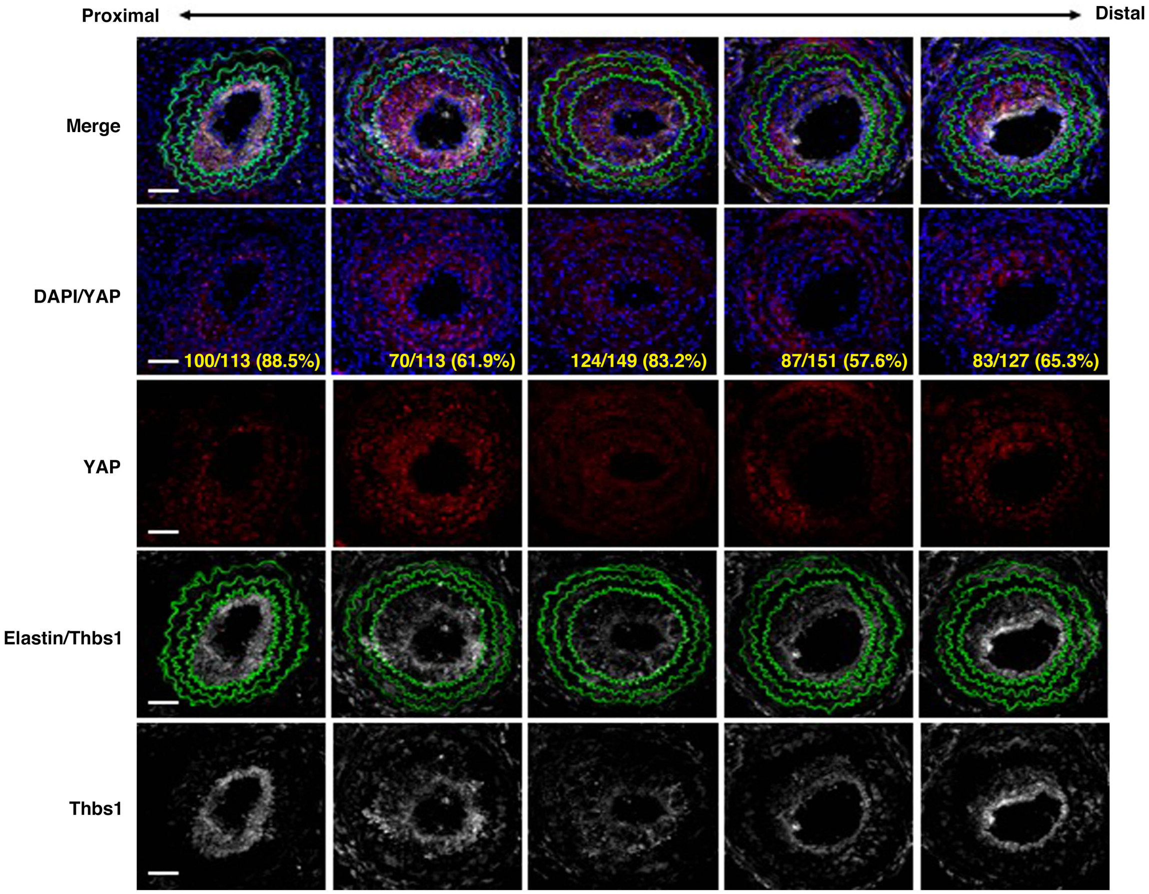


**Online Figure XVI. Neointima formation at 3 weeks after carotid artery ligation.** Representative immunostaining of cross sections of ligated arteries with YAP (red) and Thbs1 (white) at 3 weeks after ligation in WT mice (n=4). DAPI (blue) and auto-fluorescence of elastin (green). Images shows are shown in proximal to distal direction from the ligature point. Bars are 50 μm. Quantification of co-localization with YAP and DAPI using Imaris colocalization software were indicated by yellow (number of nuclear YAP / total number of nucleus).

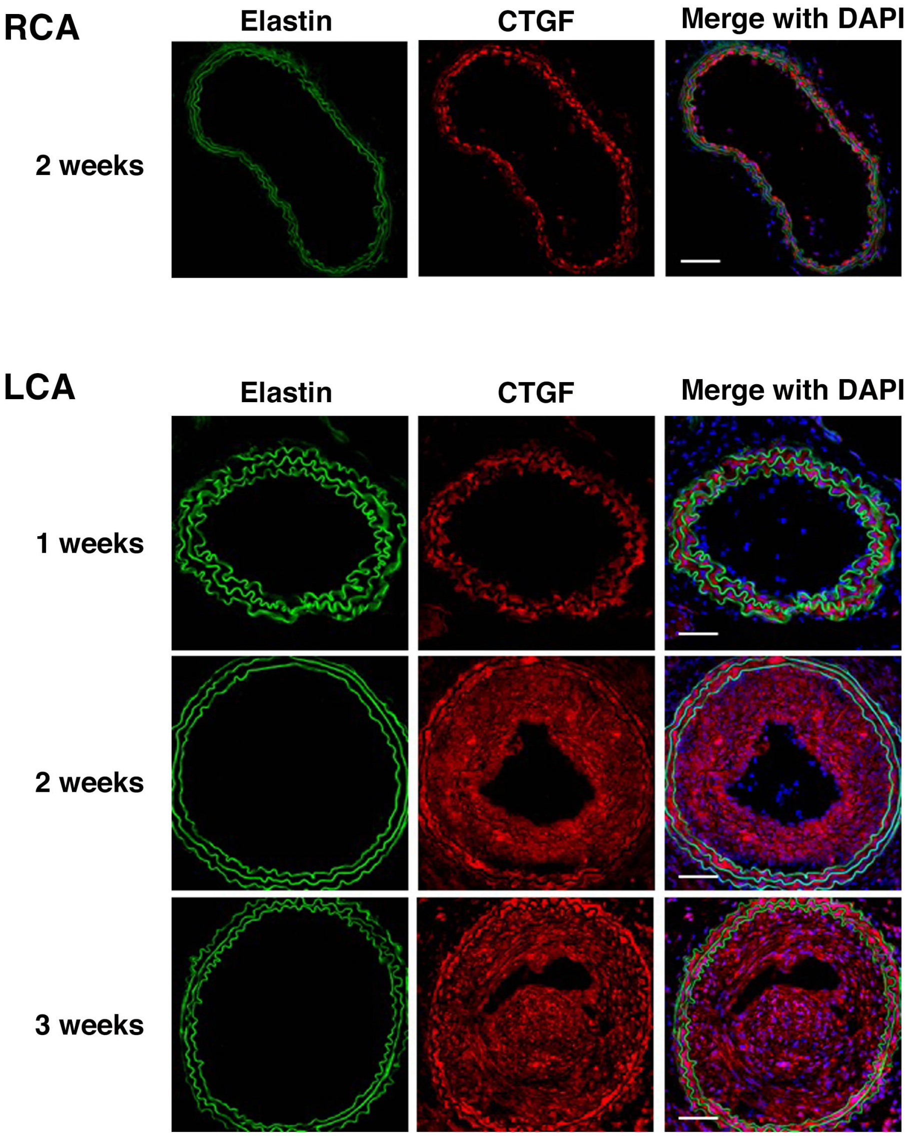


**Online Figure XVII. CTGF, downstream target of YAP, is expressed in neointima after 2 weeks of carotid artery ligation.** Representative immunostaining of cross sections of right carotid artery (RCA) and left carotid artery (LCA) with CTGF (red) after ligation in WT mice (n=3). DAPI (blue) and auto-fluorescence of elastin (green). Bars are 50 μm.

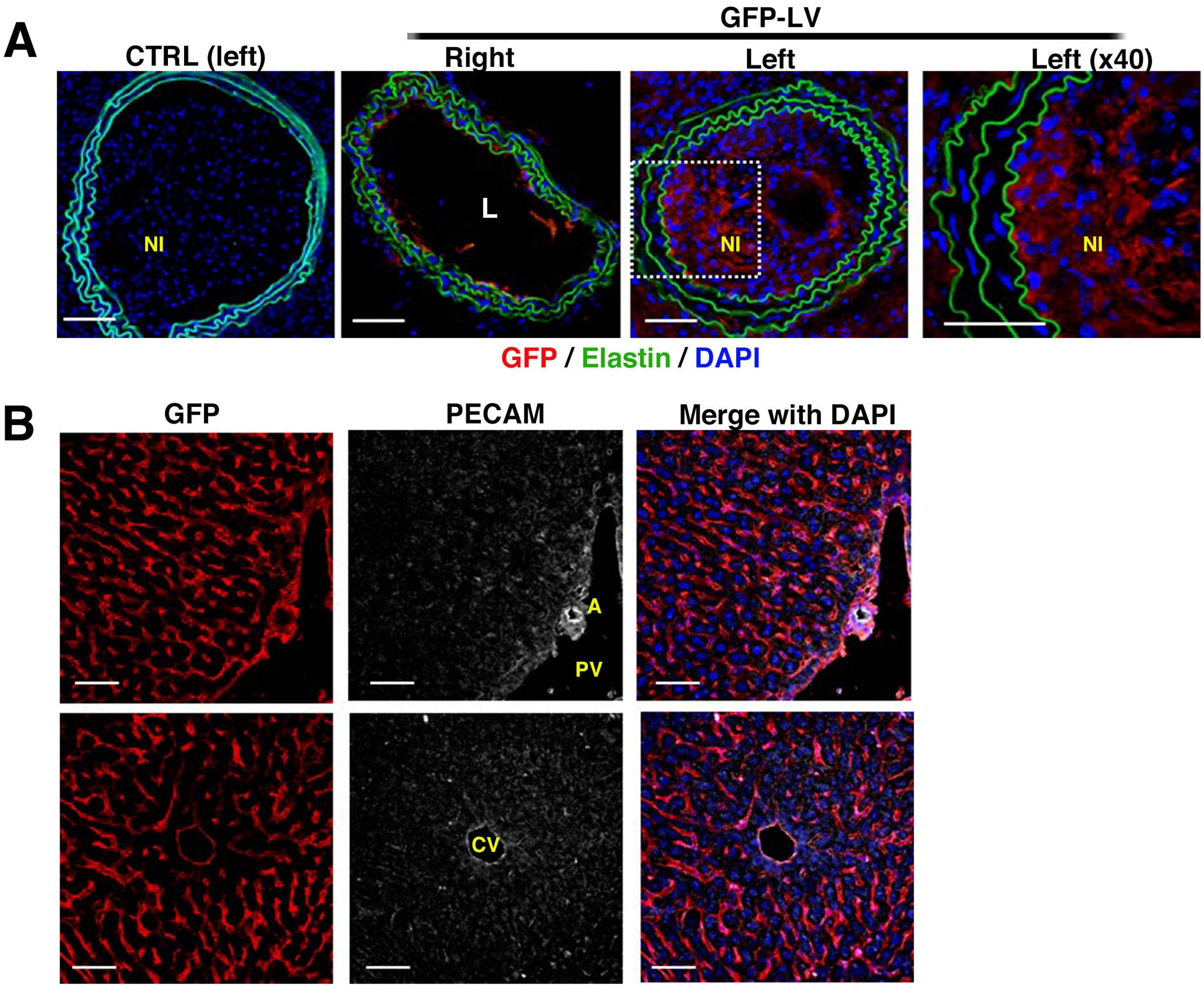


**Online Figure XVIII. Lenti-virus (LV) were delivered from tail vein to neointima of carotid artery.** GFP-LV were injected from tail vein 2 weeks after left carotid ligation. **A,** Cross sections of carotid artery at 3 weeks post injury in WT mice (CTRL, n=4), GFP-LV-treated WT mice (n=3). Immunostaining with GFP (red). DAPI (blue) and auto-fluorescence of elastin (green). Note neointima (NI) formation in left carotid artery. Bars are 50 μm. **B,** Liver sections at 3 weeks post injury in GFP-LV-treated WT mice (n=3). Immunostaining with GFP (red), PECAM (white) and DAPI (blue). A: artery, PV: portal vein, CV: central vein. Bars are 50 μm.

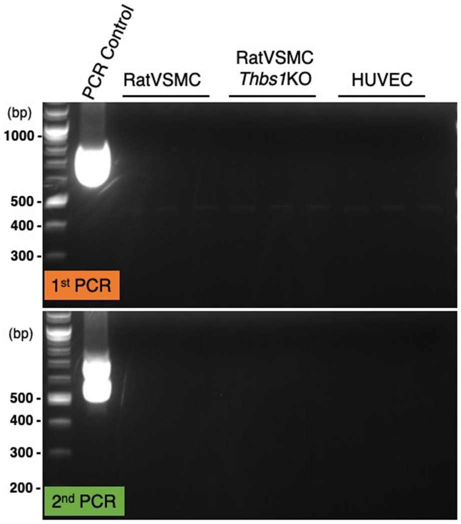


| | 1st PCR (bp) | 2nd PCR (bp) |
|---|---|---|
| *M. hyopneumoniae* | 681 | 237 |
| *M. neurolyticum* | 501 | 196 |
| *M. fermentans* | 491 | 195 |
| *M. pulmonis* | 477 | 189 |
| *M. hyorhinis* | 448 | 211 |
| *M. orale* | 423 | 179 |
| *M. capricolum* | 415 | 179 |
| *M. arthritidis* | 408 | 157 |
| *M. salivarium* | 403 | 151 |
| *M. hominis* | 370, 369 | 147, 148 |
| *M. arginini* | 369 | 145 |
| *U. urealyticum* | 482, 481 | 154 |

**Online Figure XIX.** Cell lines were tested for mycoplasma contamination. 1st PCR and nested 2nd PCR results are shown (n=3). Control template was used as positive control for PCR. The table shows each PCR product size (bp) for *Mycoplasma.*

**Online Table I. Antibodies used in this study**

**Western blot analysis**

| Antibody | Dilution | Source | Catalog number |
|---|---|---|---|
| **Thbs1** | 1:1000 | NeoMarkers | MS-421 |
| **GAPDH** | 1:1000 | Cell Signaling | # 2118 |
| **pERK1/2** | 1:500 | Cell Signaling | # 4376 |
| **ERK** | 1:1000 | Cell Signaling | # 9102 |
| **Egr1** | 1:500 | Cell Signaling | # 4154 |
| **pp38** | 1:500 | Cell Signaling | #4511 |
| **pFAK** | 1:500 | Cell Signaling | #3283 |
| **Integrin αv** | 1:500 | Cell Signaling | #4711 |
| **Integrin β1** | 1:500 | Cell Signaling | #4706 |
| **pYAP** | 1:500 | Cell Signaling | #4911 |
| **pLATS1 (T1079)** | 1:500 | Cell Signaling | #8654 |
| **His** | 1:1000 | MBL Life Science | D291-3 |

**Immunostaining**

| Antibody | Dilution | Source | Catalog number |
|---|---|---|---|
| **Thbs1** | 1:200 | NeoMarkers | MS-421 |
| **Rhodamine Phalloidin** | 1:500 | Cytoskeleton | # PHDR1 |
| **pPaxillin (Tyr118)** | 1:200 | Santa Cruz Biotechnology | sc-14036 |
| **Integrin αv (N-19)** | 1:200 | Santa Cruz Biotechnology | sc-6616 |
| **Integrin β1 (N-20)** | 1:200 | Santa Cruz Biotechnology | sc-6622 |
| **Integrin β3 (H-96)** | 1:200 | Santa Cruz Biotechnology | sc-14009 |
| **YAP** | 1:200 | Cell Signaling | #14074 |
| **Vinculin** | 1:200 | Invitrogen | #700062 |
| **c-Myc** | 1:200 | Santa Cruz Biotechnology | sc-40 |
| **CD36 (H-300)** | 1:200 | Santa Cruz Biotechnology | sc-9154 |
| **CD47 (H-100)** | 1:200 | Santa Cruz Biotechnology | sc-25773 |
| **VE-cadherin (F-8)** | 1:200 | Santa Cruz Biotechnology | sc-9989 |
| **CTGF** | 1:300 | Bio Vision | 5553R |
| **α-SMA (1A4)** | 1:200 | Sigma-Aldrich | A2547 |
| **PECAM (M-20)** | 1:200 | Santa Cruz Biotechnology | Sc-1506 |
| **GFP** | 1:500 | Millipore | MAB2510 |

**Online Table II. qPCR primer sequences**

| | |
|---|---|
| rat *Gapdh* | forward 5'-AATGTATCCGTTGTGGATCTGAC-3' |
| | reverse 5'-TCTCTTGCTCTCAGTATCCTTGC -3' |
| rat *Itgav* | forward 5'- TGAGGATCTCTTCAACTCTACGC -3' |
| | reverse 5'- AAATACCAACACGGCCAGTAAC -3' |
| rat *Itgβ1* | forward 5'- GAGACTCCAGACTGTCCTACTGG -3' |
| | reverse 5'- CACTTGGCATTCATTTTCTCCT -3' |
| rat *Itgβ3* | forward 5'- AGGCCTCAAGTCTTGTGTGG -3' |
| | reverse 5'- TAGTGAAAGATTGCTCCTTCTGC -3' |
| rat *Fn1* | forward 5'- GAGTACATTGTCACCATCATTGC -3' |
| | reverse 5'- TGATTCTGTAATAGCGCACAGAG -3' |
| rat A*cta2* | forward 5'- GTCGCCATCAGGAACCTCGASAA -3' |
| | reverse 5'- CACCAGCAAAGCCCGCCTTACA -3' |
| rat *Myh11* | forward 5'- AAGACAGCAGCATCACGGGGA -3' |
| | reverse 5'- TGCCAAAGCGGGAGGAGTTGTC -3' |
| rat *Fbn1* | forward 5'- CCGGATGGAAAACCTTACCT -3' |
| | reverse 5'- GCAAGTGCACATGTTTGGTC -3' |
| rat *Eln* | forward 5'- GAGTTAGGGTCCCAGGTGCT -3' |
| | reverse 5'- CAGCTTTGGCAGCAGCTT -3' |
| rat *Lox* | forward 5'- TTGAGTCCCGGATGTTATGA -3' |
| | reverse 5'- CCAGGTAGCTGGGGTTTACA -3' |
| rat *Cpeb3* | forward 5'- TCTCCAATCAGCAGAGCTACATT -3' |
| | reverse 5'- TAGTACTGCAGACAGGTGACGTT -3' |
| rat *Tet3* | forward 5'- CGTCTGGGACTGAAGGAAGG -3' |
| | reverse 5'- CATCTTGTACAGGGGTAGCACAT -3' |
